# Complete assembly of *Escherichia coli* ST131 genomes using long reads demonstrates antibiotic resistance gene variation within diverse plasmid and chromosomal contexts

**DOI:** 10.1101/558635

**Authors:** Arun Gonzales Decano, Catherine Ludden, Theresa Feltwell, Kim Judge, Julian Parkhill, Tim Downing

## Abstract

The incidence of infections caused by extraintestinal *Escherichia coli* (ExPEC) is rising globally, which is a major public health concern. ExPEC strains that are resistant to antimicrobials have been associated with excess mortality, prolonged hospital stays and higher healthcare costs. *E. coli* ST131 is a major ExPEC clonal group worldwide with variable plasmid composition, and has an array of genes enabling antimicrobial resistance (AMR). ST131 isolates frequently encode the AMR genes *bla*_CTX-M-14/15/27_, which are often rearranged, amplified and translocated by mobile genetic elements (MGEs). Short DNA reads do not fully resolve the architecture of repetitive elements on plasmids to allow MGE structures encoding *bla*_CTX-M_ genes to be fully determined. Here, we performed long read sequencing to decipher the genome structures of six *E. coli* ST131 isolated from six patients. Most long read assemblies generated entire chromosomes and plasmids as single contigs, contrasting with more fragmented assemblies created with short reads alone. The long read assemblies highlighted diverse accessory genomes with *bla*_CTX-M-15_, *bla*_CTX-M-14_ and *bla*_CTX-M-27_ genes identified in three, one and one isolates, respectively. One sample had no *bla*_CTX-M_ gene. Two samples had chromosomal *bla*_CTX-M-14_ and *bla*_CTX-M-15_ genes, and the latter was at three distinct locations, likely transposed by the adjacent MGEs: IS*Ecp1*, IS*903B* and Tn*2*. This study showed that AMR genes exist in multiple different chromosomal and plasmid contexts even between closely-related isolates within a clonal group such as *E. coli* ST131.

**Importance:** Drug-resistant bacteria are a major cause of illness worldwide and a specific subtype called *Escherichia coli* ST131 cause a significant amount of these infections. ST131 become resistant to treatments by modifying their DNA and by transferring genes among one another via large packages of genes called plasmids, like a game of pass-the-parcel. Tackling infections more effectively requires a better understanding of what plasmids are being exchanged and their exact contents. To achieve this, we applied new high-resolution DNA sequencing technology to six ST131 samples from infected patients and compared the output to an existing approach. A combination of methods shows that drug-resistance genes on plasmids are highly mobile because they can jump into ST131’s chromosomes. We found that the plasmids are very elastic and undergo extensive rearrangements even in closely related samples. This application of DNA sequencing technologies illustrates at a new level the highly dynamic nature of ST131 genomes.

## Introduction

Reported cases of bloodstream and urinary tract infections caused by extraintestinal pathogenic *Escherichia coli* (ExPEC) are increasing globally at an alarming rate [1]. As a key source of ExPEC isolates worldwide, *E. coli* sequence type 131 (ST131) is regarded as a serious threat to public health, given its high level of antimicrobial resistance (AMR), as well as the broad spectrum of infections it causes in community and hospital settings [2,3].

*E. coli* ST131 is virulent [4] and has an expansive range of virulence factors [5,6], especially those linked to uropathogenic *E. coli* (UPEC) [3,7,8]. AMR and virulence genes allow ST131 to adapt to drug selection pressure and to survive in extraintestinal niches, and are often encoded on mobile genetic elements (MGEs) [9], which means the exact set of virulence and AMR genes in a single ST131 isolate may vary [8,10]. ST131 encodes a range of extended-spectrum β-lactamases (ESBLs) that hydrolyse third-line drugs including cephalosporins, the most common of which encode cefotaximase *bla*_CTX-M-15_. Within ST131, clade C2 has more AMR genes than other clades and is typically *bla*_CTX-M-15_-positive, differentiating it from clade C1 that can be *bla*_CTX-M-14_ or *bla*_CTX-M-27_-positive [3,8].

Most ST131 AMR genes are reported to be encoded on plasmids: circular self-replicating double-stranded DNA molecules that constitute part of the bacterial accessory genome [11–13]. Plasmids can reduce bacterial cell fitness, but a number of post-segregation killing and stable plasmid inheritance mechanisms allow the stable maintenance of IncF plasmids in ST131 [14–16]. The chromosomal integration of plasmid genes is most commonly facilitated by transposons, which can ensure acquisition and conservation of such elements if there is no subsequent local recombination [17–18].

Identifying plasmid conjugation, recombination and transposition could have value in tracking AMR genes associated with disease outbreaks and antibiotic treatment failures. Plasmids may be classified using incompatibility (Inc), relaxase (MOB) and mating pair formation system typing [19], but difficulties in plasmid genetic analysis and reconstruction arise with short read data due to rearrangements driven by recombination, dense arrays of repetitive elements including transposable elements (TEs), changes in gene copy numbers, and high sequence variation. Methods using short reads alone may fail to detect genomic segments exchanged between plasmids and the chromosome, limiting evaluation of the core and accessory genomes.

Whole genome sequencing has provided a high resolution of the genomic epidemiology of ST131 and plasmid-mediated AMR outbreaks [20]. However, short reads alone are insufficient to resolve plasmids that often have numerous small MGEs of ~1 kb or less in size, e.g. TEs and insertion sequences (ISs) [21]. Complex transposable units (TUs) consisting of multiple TEs or ISs can mobilise AMR genes by transposition, and this can sometimes be followed by recombination within the TU between one of the inverted repeats (IRs) flanking the TE and the IR of another local TE or an adjacent homologous sequence, resulting in different TU structures, locations and copy numbers. At present, the exact resolution of complex structural rearrangements of repetitive TUs containing AMR genes may be impossible with short reads [22]. Consequently, plasmid assembly is a challenge requiring accurate long reads and sufficient coverage to distinguish between independent plasmids with regions of sequence identity [21,23].

Long reads, such as those generated using Oxford Nanopore Technologies (ONT) or Pacific Biosciences platforms can provide a solution to this plasmid assembly problem [24–26]. Here, we sequenced six ST131 using the ONT GridION X5 platform. Using the resulting high-coverage sequence data, we reconstructed and annotated the plasmids and chromosomal regions carrying *bla*_CTX-M_ genes, as well as their genetic context and copy numbers.

## Methods

### Sample collection

Six ESBL-producing *E. coli* ST131 clinical strains were isolated in June-October 2015 from patients at Addenbrooke’s Hospital, Cambridge, as part of a study on antibiotic resistance (Supplementary Table 1). Five samples were from faeces, and one was from blood. These were short-read sequenced in a multiplex run on an Illumina HiSeq 2500 platform and processed as previously outlined [27].

### High molecular weight DNA extraction

Frozen stocks of the six isolates were streaked onto LB agar plates and grown overnight at 37°C. Single colonies were subcultured onto LB agar plates and incubated overnight at 37°C. DNA was extracted using a Lucigen Masterpure Complete DNA and RNA Purification kit. For each sample, a swab was used to sweep half a plate of pure colonies, and suspended in 1x phosphate buffer solution (PBS). Samples were processed according to the manufacturer’s instructions, with elution in 70ul of Nuclease Free water. Pipetting was minimised to reduce shearing of the DNA prior to sequencing.

### Oxford Nanopore library preparation and sequencing

DNA was quantified using a Quant-iT™ HS (High Sensitivity) kit (Invitrogen). DNA purity was checked using a Nanodrop (ThermoFisher) and fragment size was confirmed by FEMTO Pulse (Nano Life Quest). The sequencing libraries were prepared using 1 μg DNA per sample and ligation sequencing kit 1D SQK-LSK109 with the barcoding extension kit EXP-NPB104 according to ONT protocols. The samples were combined using equimolar pooling and loaded onto a single 9.4.1 MIN-106 flow cell and sequenced on the GridION X5 platform under standard conditions.

### Illumina library preparation and sequencing

The short reads used in this study were created as follows: bacterial genomic DNA was extracted using the QIAxtractor (Qiagen, Valencia, CA, USA) according to the manufacturer’s instructions. Library preparation was conducted according to the Illumina protocol and sequenced (96-plex) on an Illumina HiSeq 2500 platform (Illumina, San Diego, CA, USA) using 100 bp paired-end reads.

### Oxford Nanopore base-calling and adapter trimming

The resulting fast5 read files were transferred to a separate Linux server 4.4.0 (Ubuntu 16.04.4) for analysis. Basecalling was performed during the GridION run using ONT’s Guppy v0.5.1 and the resulting fast5 from the initial run was converted to fastq format with Albacore v2.0 (ONT). The statistical data of the sequencing run was processed with MinIONQC v1.3.5 [28] based on the default Q score cut-off of seven. Adapters and chimeric reads were removed from fastq files using Porechop v0.2.4 [29] with demultiplex settings (Supplementary Figure 1). Standard outputs were saved as log files and were then parsed. The quality of the final fastq files was assessed using FastQC v0.11.8 (https://www.bioinformatics.babraham.ac.uk/projects/fastqc/) and MultiQC v1.4 [30].

### Genome assembly and improvement

We assembled the genomes using the conservative, normal and bold modes of the long read-only assembly pipeline in Unicycler v4.6. Previous work has suggested that Unicycler outperforms alternatives [21] that struggle to resolve plasmids [31]. This workflow included the assembly polisher, Racon, which ran iteratively to minimise error rates of called bases [29]. For comparison, short read-only and hybrid assemblies were also created using Unicycler v4.6. Briefly, during short read-only assembly, Unicycler v4.6 employed SPAdes v3.12 to assemble short reads then used Pilon v1.22 to polish the assembly. In hybrid assemblies, Unicycler v4.6 used Miniasm to piece long reads together first and applied SPAdes v3.12 to incorporate short reads and bridge gaps. Pilon was run 3-10 times for short read assemblies and 5-10 times for hybrid ones, until no further changes were required to achieve the most contiguous and completed genome assemblies. The average number of changes by Pilon was 74.3, 100.2 and 125.3 for short read assemblies, and 234.5, 257.7 and 377.0 across conservative, normal and bold modes (respectively).

### Genome assembly assessment and error rate quantification

The quality of resulting assemblies was assessed using Quast 3.0 [32] according to the total assembly length, number of contigs, N50, GC content and degree of replicon circularization. Assembly graphs were visualized with Bandage [33]. The resulting contigs in each assembly were classified as chromosomal or plasmid using machine learning algorithms implemented in mlplasmids [22]. Genome completeness was examined using the numbers of single-copy universal orthologous genes using Benchmarking Universal Single-Copy Orthologs (BUSCO) v3 with the gammaproteobacteria_odb9 database [34].

### Read depth estimation

The read depth of each replicon was estimated by aligning the short Illumina and long Oxford Nanopore reads to the completed genomes using Smalt v0.7.6 and BWA-MEM v0.7.17 (with the flag –x ont2d for ONT reads), respectively. SAMtools v1.7 was used to process the SAM files to BAM format, remove duplicates, and identify the coverage at each base of each assembly. The median value for each replicon was noted and was normalized using the median chromosomal depth of the same assembly.

### Genome annotation

The genomes were annotated using Prokka v1.13.3 [35]. *Bla*_CTX-M_ alleles and their contexts were detected using the Multiple Antibiotic Resistance Annotator (MARA) [36] and by aligning the assemblies against the Comprehensive Antibiotic Resistance Database (CARD v3.0) to screen for matches with 100% ID only. Information on the detected AMR features and MGEs are retrieved from Galileo AMR (https://galileoamr.arcbio.com/mara/feature/list). Plasmid identification and typing was carried out using PlasmidFinder v2.0 [37]. The plasmid-derived contigs from the assembled genomes were compared using BLAST v2.6.0 using a database of 10,892 complete plasmids [38]. Their homology and annotation was visualised using EasyFig v2.2.2 [39].

### Phylogenetic analysis

To provide a phylogenetic context for these six isolates, the short Illumina reads of 63 from [8] and 56 from [38] published ST131 short read libraries were cleaned and trimmed using Fastp v0.12.3 [41], as were the six isolates’ short read libraries from this study. These 125 libraries were *de novo* assembled with Unicycler v4.6 using NCTC13441 as a reference and annotated using Prokka. The 126 genomes were processed using Roary v3.11.2 [42] with a 95% BLAST v2.6.0 identity threshold to create a core genome alignment containing 4,457 SNPs using MAFFT v7.310 [43] spanning 3,250,343 bases and 3,350 genes of the NCTC13441 chromosome (a length similar to [20]). This core genome was used to construct a maximum likelihood phylogeny using RAxML v8.2.11 with the GTR model with gamma rate heterogeneity [44]. Clade classification of the six isolates was based on published ST131 phylogenetic analysis [8] with associated classification and *bla*_CTX-M_ allele data from [8] and [40].

## Results

### Oxford Nanopore long read quality control and filtering

High molecular weight DNA from six *E. coli* ST131 isolates was sequenced using long Oxford Nanopore reads and short Illumina reads to assemble their genomes allowing for plasmid reconstruction and resolution of AMR genes, MGEs and associated rearrangements. The ONT GridION X5 sequencing generated 8.9 Gbases in total across 1,406,087 reads (mean length of 6.3 Kb, Table 1). The number of reads generated per hour, total yield of bases over time, read length distribution, and read Q score distribution were examined (Supplementary Figure 2). Half of the bases with Q ≥ 7 were on reads of 18 Kb or longer (Supplementary Figure 3). These metrics indicated sufficient GridION data in terms of quantity and quality. Initial screening removed reads with Q < 7, leaving 1,142,067 reads with 8.2 Gbases with a mean Q score of 10.2 and a mean length of 7.2 Kb (Table 1) for analysis. This included 81 reads longer than 100 Kb, including one of 155,312 bases. This corresponded to 257-fold theoretical coverage for six 5.3 Mb genomes.

**Table 1.**
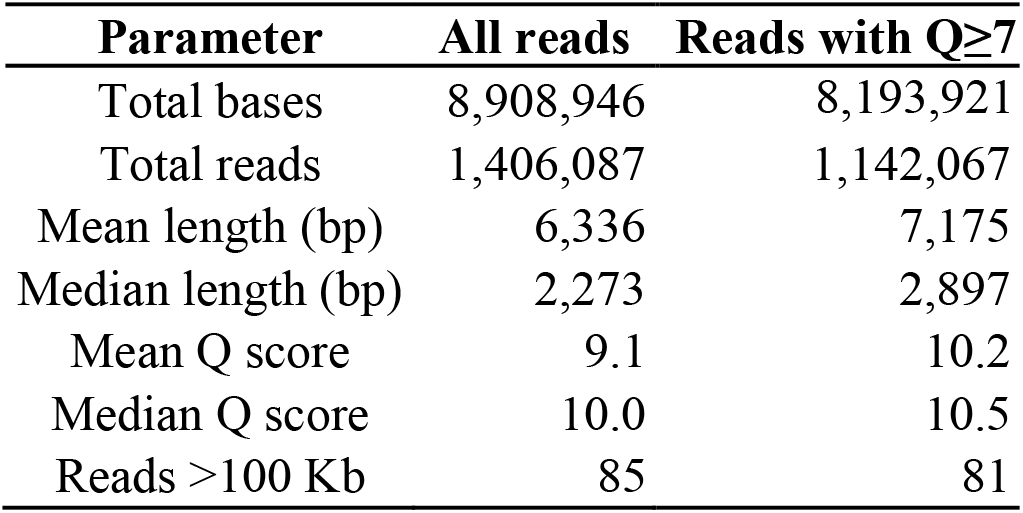
Quality parameters indicated high-quality read libraries for the six ST131 samples from GridION X5 sequence data. A total of 264,020 of low-quality reads (with Q<7) totalling 715,024,800 bases were excluded.

The initial number of reads per library ranged from 127,118 to 510,253 and these were filtered using a series of steps to ensure that the reads used for each of the six assemblies had high quality. Bases were successfully called at an average of 97.9% of reads (Table 2). Identifying the consensus demultiplexed, duplicate-free and adapter-free reads from Porechop v0.2.4 eliminated a further 2.9% of the basecalled reads, yielding 120,123 to 487,482 reads per library (Table 2).

**Table 2.**
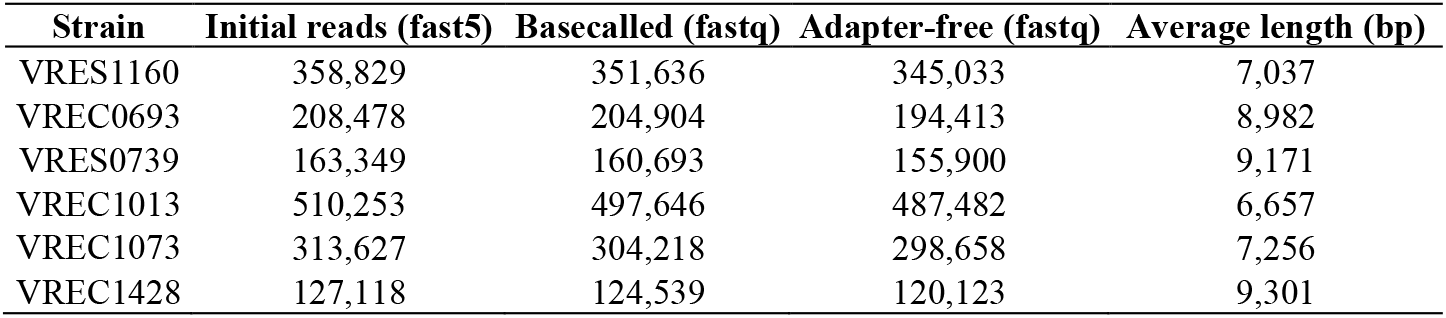
Number of reads generated from GridION X5 sequencing data per library that passed filtering during basecalling with Albacore v2.0 and those that were adapter-free (using Porechop v0.2.4). The latter totalling 1,601,609 reads were used for downstream analyses. 80,045 reads were excluded during basecalling or adapter-trimming.

| Strain | Initial reads (fast5) | Basecalled (fastq) | Adapter-free (fastq) | Average length (bp) |
| --- | --- | --- | --- | --- |
| VRES1160 | 358,829 | 351,636 | 345,033 | 7,037 |
| VREC0693 | 208,478 | 204,904 | 194,413 | 8,982 |
| VRES0739 | 163,349 | 160,693 | 155,900 | 9,171 |
| VREC1013 | 510,253 | 497,646 | 487,482 | 6,657 |
| VREC1073 | 313,627 | 304,218 | 298,658 | 7,256 |
| VREC1428 | 127,118 | 124,539 | 120,123 | 9,301 |

### Long read genome assembly illuminates highly diverse accessory genomes

All six genome assemblies produced chromosomes of 4.81-5.38 Mb with differing numbers of plasmids with lengths spanning 4-156 Kb (Supplementary Figure 4; Table 3). The numbers of contigs produced by long read assemblies of two samples (VREC0693, VRES0739) corresponded exactly to the chromosome and plasmids. The others had either one (VREC1073, VRES1160, VREC1013) or two (VREC1428) additional chromosomal contigs (Supplementary Table 2).

**Table 3.** Total size of assemblies, chromosomes and plasmids found in each strain based on their optimal whole genome assemblies using the GridION X5 long reads. Each assembly had seven or less contigs, and in three cases no fewer contigs were possible, consistent with full genome assembly (for VREC0693, VRES0739 and VREC1073). The optimal assembly with Unicycler used long reads alone (in bold mode), with exception of VREC1013, where a hybrid combining short Illumina reads with long Oxford Nanopore reads was best, with minor manual screening (Supplementary Results).

| Strain | Genome length (bp) | Number of contigs |  | N50 | Chromosome size (Mb) | Number of plasmids | Plasmid sizes (Kb) |
| --- | --- | --- | --- | --- | --- | --- | --- |
|  |  | Assembled | Minimum possible |  |  |  |  |
| VRES1160 | 5,326,801 | 6 | 5 | 5,126,679 | 5.23 | 4 | 62, 16, 5, 4 |
| VREC0693 | 5,260,741 | 3 | 3 | 5,039,909 | 5.04 | 2 | 132, 89 |
| VRES0739 | 4,806,912 | 3 | 3 | 4,797,749 | 4.81 | 2 | 5, 4 |
| VREC1013 | 5,223,433 | 3 | 2 | 3,699,451 | 5.14 | 1 | 90 |
| VREC1073 | 5,539,158 | 3 | 2 | 5,286,804 | 5.38 | 1 | 156 |
| VREC1428 | 5,236,419 | 7 | 5 | 4,924,536 | 5.13 | 4 | 92, 5, 5, 4 |

Contigs were classified as chromosomal or plasmid-derived using mlplasmids given a probability threshold of 60% [22], with further screening for plasmid-related gene content using MARA, CARD and PlasmidFinder (Supplementary Table 2). The largest plasmid was a 156.3 Kb IncFIA one in VREC1073, its sole plasmid. VREC1428 and VRES1160 had 92.8 and 61.9 Kb IncFIA plasmids, respectively, along with three small Col plasmids each (Table 3). VREC0693 had a 132.0 Kb IncFIB plasmid and an 88.8 Kb IncB plasmid - IncB plasmids have the same Rep domains as IncFII plasmids [45]. VREC3013 had one 89.9 Kb IncFII plasmid. VRES0739 alone had no large plasmid, which was verified with the short read data.

By mapping the long reads to the optimal assemblies, the read coverage of each chromosome and plasmid was estimated (Supplementary Table 2). Each chromosome had between 126- and 310-fold median coverage, and the median coverage levels of large plasmids ranged from 85- to 282-fold, except for VREC1013’s IncFII plasmid that had 1,015-fold coverage and a normalized depth of 3.3-fold. The normalised depth of plasmids compared to chromosomes suggested some cells in VREC1428 and VREC1073 may have lost their IncFIA plasmid, and the same for VREC0693 and its IncFIB plasmid. However, the IncFIA plasmid in VRES1160 and the IncB plasmid in VREC0693 had higher than expected copy numbers (by 9% after normalisation), potentially indicating stable plasmid retention.

Across five assemblies in the Unicycler normal mode, the median indel error rates for short reads and hybrid assemblies were similar (0.21 and 0.28 per 100 Kb, respectively), but was much higher for long read assemblies (265.0 per 100 Kb, Supplementary Table 3). Likewise, the median mismatch error rates for short reads and hybrid assemblies were comparable (4.25 and 2.28 per 100 Kb, respectively), but was much higher for long read assemblies (332.8 per 100 Kb, Supplementary Table 3). These rates excluded VREC1073, for which some Quast metrics were zero values. Similarly, the recovery of conserved BUSCO genes was far higher for hybrid assemblies (>99.5%) than for long read ones (>82.3%).

### The dynamic locations and genomic contexts of *bla*_CTX-M_ genes in long read assemblies

The optimised assemblies provided an improved view of the genomic context of each *bla*_CTX-M_ allele, whose effectiveness as a marker for ST131 clade classification and origin [8] we explored here. The deeper resolution of genome architecture revealed surprising differences in *bla*_CTX-M_ gene context (Figure 1; Supplementary Table 2), including the discovery of chromosomal *bla*_CTX-M_ genes in VREC0693 (three copies of *bla*_CTX-M-15_) and VREC1073 (one copy of *bla*_CTX-M-14_). All *bla*_CTX-M_ genes were complete (876 bp) with adjacent IS*Ecp1* (1,658 bp with flanking IRs of 14-16 bp) and Tn*2* (5.8 Kb) elements: IS*Ecp1* and Tn*2* can transpose *bla*_CTX-M_ and other ESBL genes [46–47]. The VRES0739 genome did not contain any region homologous to *bla*_CTX-M_, most likely because it had lost an IncF plasmid, unlike the other isolates.

**Figure 1.**
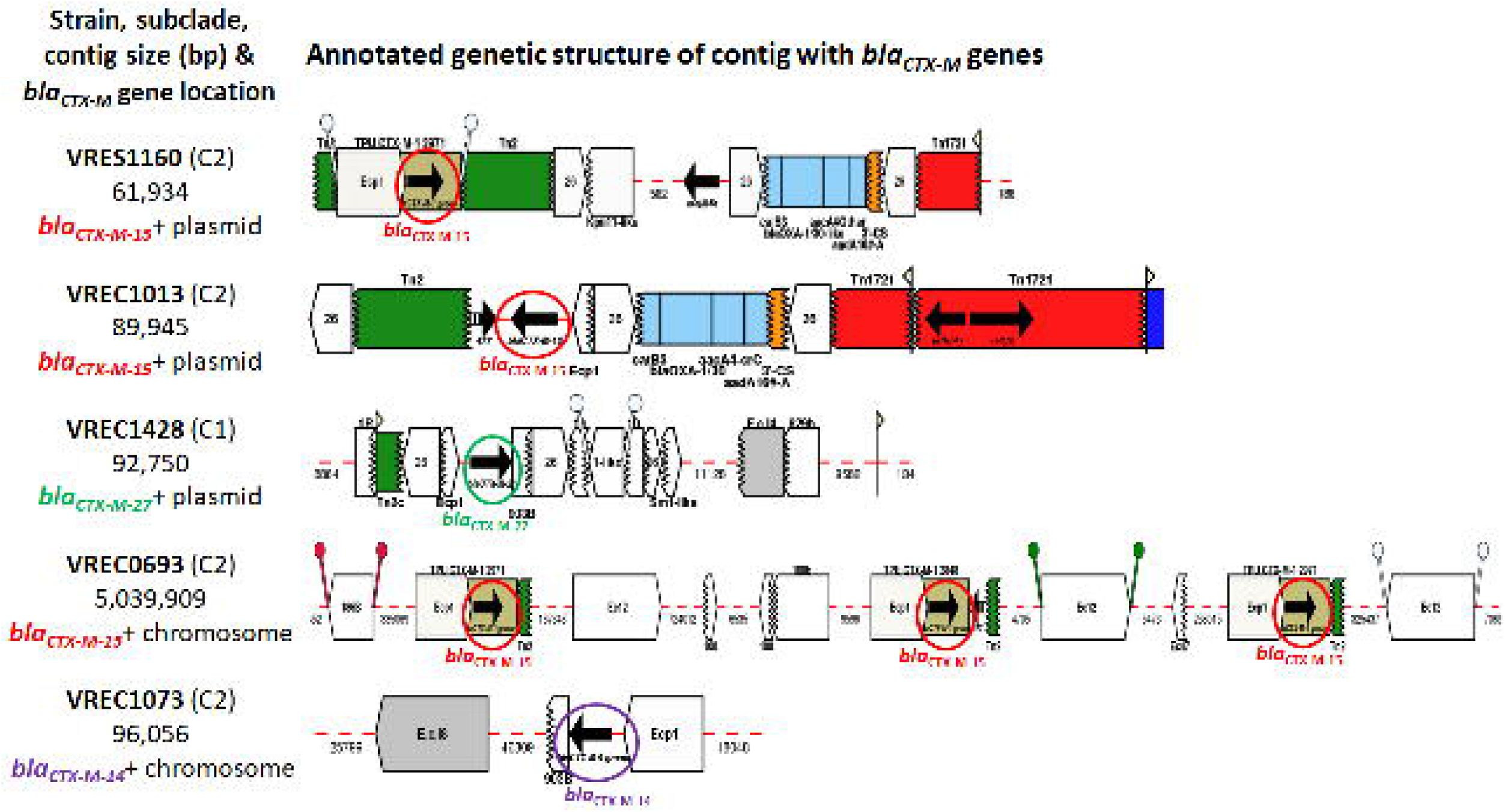
Two of the ST131’s *bla*_CTX-M_ genes were on chromosomal contigs (VREC0693 and VREC1073). VRES1160 and VREC1013 had IncFIA and IncFII plasmids, respectively, both of which had *bla*_CTX-M-15_ genes. VREC1428 had an IncFIA plasmid with *bla*_CTX-M-27_ gene. VRES0739 is not shown because it was *bla*_CTX-M_-negative and had no large plasmid. The contigs were classified as chromosomal or plasmid-derived by mlplasmids so that the *bla*_CTX-M_ genes and their genetic flanking context could be examined. Annotation was derived from Galileo™ AMR based on the Multiple Antibiotic Resistance Annotator (MARA) and database. The *bla*_CTX-M_ variants are labelled and encircled in red (*bla*_CTX-M-15_), purple (*bla*_CTX-M-14_) or green (*bla*_CTX-M-27_).

VRES1160, VREC0693 and VREC1013 all had *bla*_CTX-M-15_ genes linked to isoforms of IS*Ecp1*, IS*26* and Tn*2*, implicating them in driving transposition of the TU (Supplementary Figure 5). Each was similar to the ST131 clade C2 *ISEcp1-bla*_*CTX*-M-15_-orf477**Δ** TU [8,48] but with distinct structural differences. VRES1160’s single *bla*_CTX-M-15_ gene was at 2,296 bp on its IncFIA plasmid and was flanked by IS*Ecp1* to its 5’ and Tn*2* followed by IS*26* at its 3’ end, with another Tn*2* 5’ of IS*Ecp1*. VREC0693’s three chromosomal *bla*_CTX-M-15_ genes were not tandem repeats (chromosomal locations 2,781,074, 3,696,068 and 3,970,927), but each of these TUs were identical: all had IS*Ecp1* at the 5’ ends and truncated Tn*2*s at the 3’ ends. VREC1013’s sole *bla*_CTX-M-15_ gene was located at 13,226 bp on its IncFII plasmid and was flanked by a truncated IS*Ecp1* at its 5’ end and Tn*2* at its 3’ end, with IS*26* copies 5’ and 3’ of these segments.

VREC1428’s single *bla*_CTX-M-27_ gene was on its IncFIA plasmid at position 6,018, and VREC1073’s single chromosomal *bla*_CTX-M-14_ gene started at contig position 19,746 (Supplementary Figure 5). Both their *bla*_CTX-M_ genes were flanked by a truncated IS*Ecp1* at the 5’ ends and a shortened IS*903B* at the 3’ ends suggesting that IS*Ecp1* and IS*903B* may have facilitated the transposition of the TU from the plasmid. Similar *bla*_CTX-M_ gene transposition events have been observed in ST131 clade C1 [8].

Alignment of the plasmid-derived contigs of VRES1160 (IncFIA) to VREC1013 (IncFIB) showed that the *bla*_CTX-M-15_-positive plasmids were much more similar (>83% identity) relative to VREC1428’s *bla*_CTX-M-27_-positive IncFIA plasmid, which was more distinct (Figure 2). In addition, VREC1428’s plasmid had *traI* and *traD* genes indicating conjugation machinery (Supplementary Table 4a) as well as high homology to at least one published plasmid, unlike VRES1160’s and VREC1013’s plasmids (Supplementary Table 5). This suggested that the VRES1160 and VREC1013 plasmids had homology corresponding well with *bla*_CTX-M_ gene and subclade classification, and that they were structurally different to published plasmids due to recombination.

**Figure 2.**
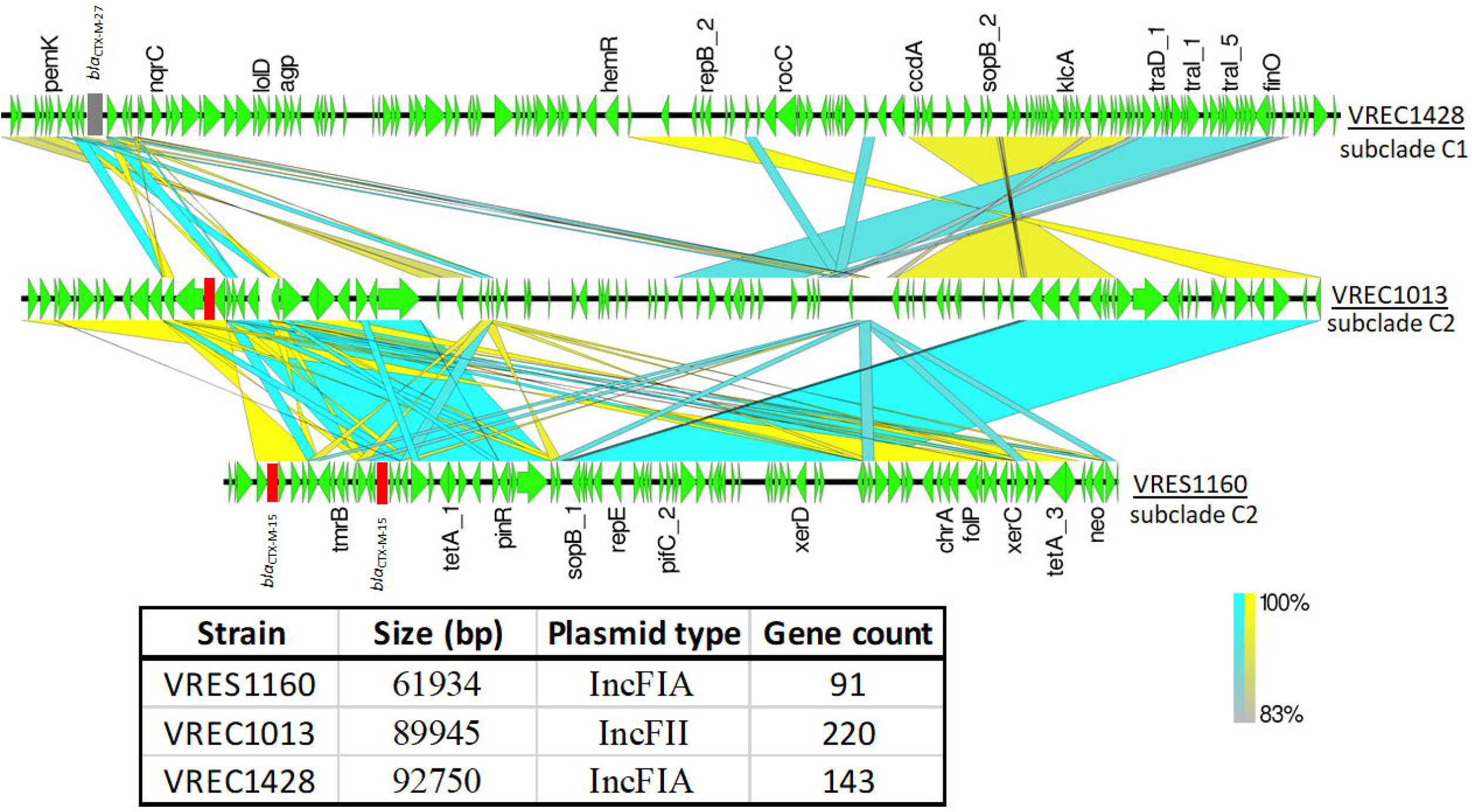
Pairwise comparisons of the three *bla*_CTX-M_-positive plasmid-associated contigs showed high sequence identity for the two from subclade C2 (VREC1013 and VRES1160) relative to one from C1 (VREC1428, top). The BLAST result was visualised with EasyFig v2.2.2 such that the middle blocks connecting regions of the contigs represent nucleotide homology: blue for homologous regions in the same direction, and yellow for inversions. Gaps or white spaces denote unique loci or regions present in a contig but not in the other. Gene models are in green with the direction of transcription shown by arrows. Genes of interest are labelled above each arrow. The *bla*_CTX-M-27_ grey (top) is in mauve and the two *bla*_CTX-M-15_ genes (middle, bottom) are in red. The table below shows the contig size, plasmid type and the number of genes per strain. The list and products of the annotated genes are in Supplementary Table 4b.

### Phylogenetic context of analysed isolates

Comparison of these six samples with 119 published ST131 [8,40] as short read assemblies scaffolded using reference genome NCTC13441 showed that all clustered in ST131 clade C (Supplementary Figure 6). There was sufficient resolution across 4,457 core genome SNPs to confidently assign them to subclades C1 (n=1) or C2 (n=5) (Figure 3). VRES1160, VREC0693, VREC1013, VRES0739 and VREC1073 clustered with C2, whereas the *bla*_CTX-M-27_-positive VREC1428 was in C1. VRES1160, VREC0693 and VREC1013 all had IncF plasmids (IncFIA, IncFIB, IncFII) and *bla*_CTX-M-15_ genes, consistent with C2 are typically *bla*_CTX-M-15_-positive, which was observed for 77% of C2 isolates here (48 out of 62). However, VREC1073 was in C2 but had an a *bla*_CTX-M-14_ gene, contradicting this pattern and was the sole *bla*_CTX-M-14_-positive C2 isolate found here. The core genomes of VRES0739 and VREC0693 were identical, implying that VRES0739 has very recently lost its (*bla*_CTX-M_-positive IncF) plasmid. The sole isolate clustering with C1 was VREC1428, which had an IncFIA plasmid with a *bla*_CTX-M-27_ gene, and so may belong to the emerging subclade C1-M27 as evidenced by the presence of prophage-like regions like M27PP1/2 [40].

**Figure 3.**
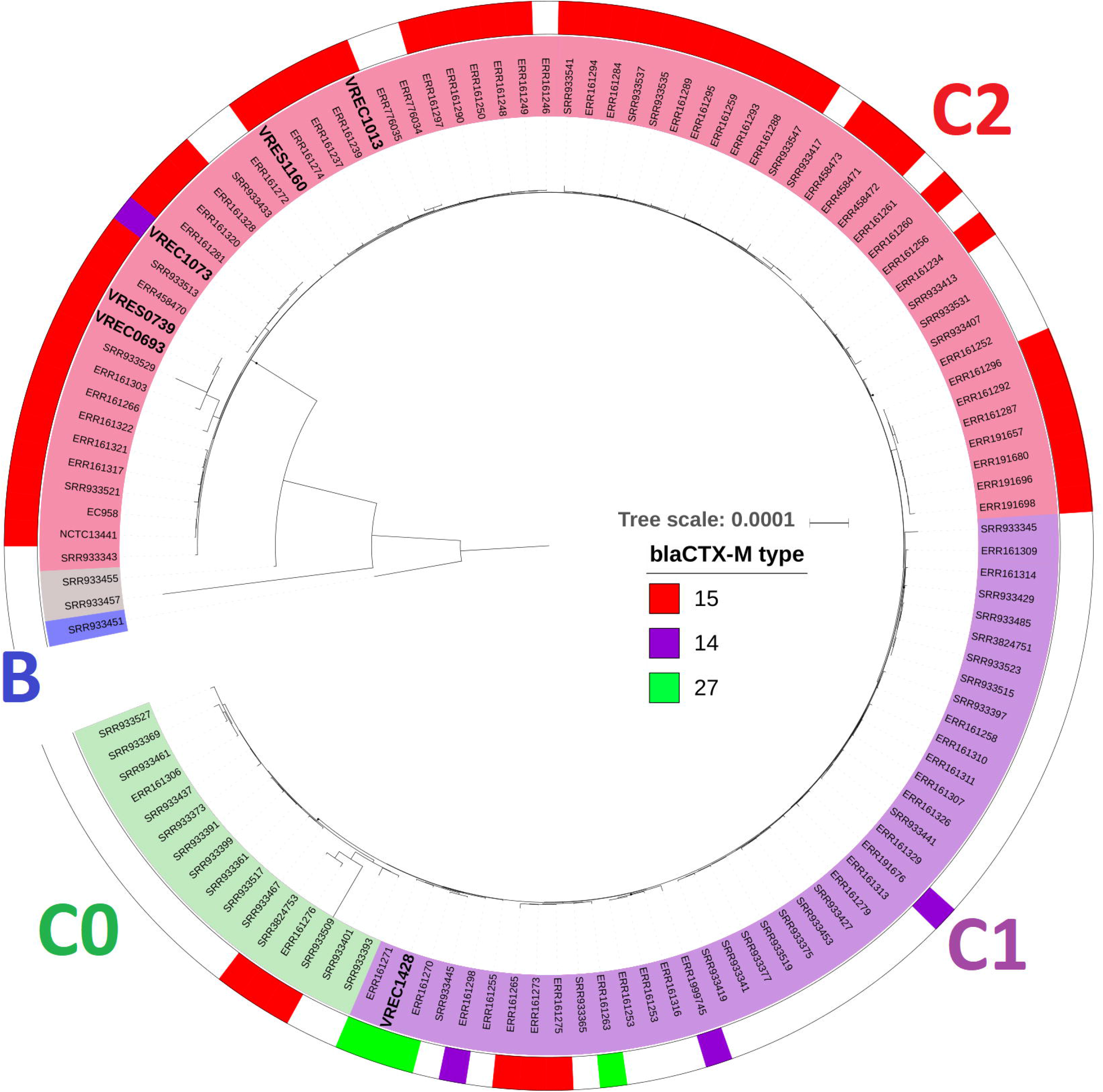
The phylogenetic context of the six ST131 genomes (names are in large bold font) showed that all except VREC1428 were in ST131 subclade C2 (red inner ring: VRES1160, VREC1073, VRES0739, VREC0693 and VREC1013). VREC1428 clustered in subclade C1 (purple inner ring). No new isolate clustered in C0 (green inner ring), B (blue inner ring) or an intermediate cluster (grey inner ring). Clade classification was based on phylogenetic analysis by [8] by including the reference NCTC13441, n=63 isolates from [8] and n=56 from [40] with associated classification and *bla*_CTX-M_ allele data. VREC1073, and VREC0693 had chromosomal *bla*_CTX-M_ genes. The outer ring shows *bla*_CTX-M-15_ (red), *bla*_CTX-M-14_ (purple) and *bla*_CTX-M-27_ alleles (green). The phylogeny was built with RAxML v8.2.11 using 4,457 SNPs from a core genome alignment generated with Roary v3.11.2 and was visualised with iTOL v4.3. Branch support was performed by 100 bootstrap replicates, and the scale bar indicates the number of substitutions per site. This mid-pointed rooted phylogeny includes reference genome isolates EC958 and NCTC13441 (both in C2).

## Discussion

Our study resolved the plasmid architecture of several recent *E. coli* ST131 isolates, allowing investigation of AMR gene location, copy number and potential transposon-driven rearrangements. This advance was facilitated by the careful DNA handling during extraction to produce large volumes of high molecular weight DNA that was pure and free from contamination, which was avoided by performing separate extraction steps to obtain small plasmids [49] overcoming a limitation for MinION sequencing [21].

The long read genome assemblies illuminated significant variation in plasmids, MGEs and *bla*_CTX-M_ gene composition that was not captured by short reads. ST131 is a globally pandemic *E. coli* clonal group [15] with diverse sources of transmission [25]. Phylogenetic comparison with published genomes [8,40] showed that five out of six isolates were from subclade C2 with one from C1. The emergence of clade C has been associated with IncF plasmids, and clade C2 with IS*Ecp1* and Tn*2* elements flanking *bla*_CTX-M-15_ genes [50–51]. Our long read assemblies showed the excision of the entire TU from the IncFIB plasmid and chromosomal integration at three distinct locations for VREC0693, and similarly chromosomal translocation of the *bla*_CTX-M-14_ gene from an IncFIA plasmid for VREC1073, mediated by IS*Ecp1* and IS*903B* based on previous work [8]. These transposition events were likely driven by recombination at adjacent transposable elements. This highlights the value of long read sequencing to resolve the location of *bla*_CTX-M_ genes and that chromosomal translocations are not rare in ST131.

A high resolution of the AMR gene structure, context and copy number is highly predictive of AMR phenotypes [52], and could lead to new insights into AMR mechanisms. However, the high indel and mismatch errors in long Oxford Nanopore reads [31,49,53–54] limits power to identify AMR isoforms that could permit genome-based antimicrobial susceptibility testing [45,55–56]. Here, the five ONT assemblies together had an average of 447-fold higher indel and 48-fold higher mismatch error rates than those for the corresponding Illumina reads, similar to previous work with MinION reads [23], and this impacted gene identification. Consequently, short reads and assembly polishing methods remain important for SNP identification and error detection until long read error rates can be reduced [57].

Our findings illustrate the diversity of AMR gene context even within recently emerged clones such as ExPEC ST131. The detection of multiple instances of chromosomally integrated ESBL genes using long reads here for *bla*_CTX-M-15_ in *E. coli* has parallels elsewhere for *bla*_OXA-181_ in *bla*_CTX-M-15_-positive *K. pneumoniae* [58] and so highlights chromosomal ESBL gene IS*Ecp1*-mediated transposition as a potential adaptive mechanism in *Enterobacteriaceae*. Further studies are needed with larger sample sizes to identify the rates and mechanisms of these dynamic changes.

## Author contributions

Conceptualization - A.D. and T.D. Methodology - A.D., T.F., K.J. and T.D. Software - A.D. Formal Analysis - A.D. Investigation - A.D., C.L., T.F. and K.J. Resources - C.L., T.F., K.J., J.P and T.D. Writing - Original Draft - A.D., T.F. and T.D. Writing - Reviewing & Editing - A.D., C.L., T.F., K.J., J.P and T.D. Visualisation - A.D. Funding Acquisition - A.D., C.L., K.J., J.P. and T.D.

## Data Summary

1. Illumina reads accession numbers: ERR2138475, ERR2138200, ERR2138591, ERR1878196, ERR2137889 and ERR1878359 in the European Nucleotide Archive (ENA) under BioProjects PRJEB21499 and PRJEB19918.
2. ONT reads ENA accession numbers: www.ebi.ac.uk/ena/data/view/PRJXXXXXX, Figshare https://doi.org/10.6084/m9.figshare.7554293.v1
3. Unicycler assemblies, Figshare https://doi.org/10.6084/m9.figshare.7560458.v2

## Ethical approval

The study protocol was approved by the National Research Ethics Service (ref:14/EE/1123), and the Cambridge University Hospitals NHS Foundation Trust Research and Development Department (ref: A093285).

## Acknowledgements

We acknowledge Anne Parle-McDermott and Emma Finlay at Dublin City University (DCU, Ireland) for guidance on DNA extraction protocols, and also Emma Betteridge, Karen Oliver and the Long Read sequencing and data teams at the Wellcome Sanger Institute (U.K.) for their assistance with sequencing.

## Conflicts of interest

JP is a consultant to Next Gen Diagnostics Llc.

## Funding information

This project was funded by a Dublin City University (DCU) O’Hare Ph.D. Fellowship, a DCU Enhancing Performance grant, a DCU Orla Benson Memorial Scholarship grant, a DCU Advanced Research Computing Centre for Complex Systems Modelling (ARC-SYM) grant, and by the Health Innovation Challenge Fund (WT098600, HICF-T5-342), a parallel funding partnership between the Department of Health and Wellcome Trust. The views expressed in this publication are those of the author(s) and not necessarily those of the Department of Health or Wellcome Trust. Catherine Ludden is a Wellcome Trust Sir Henry Wellcome Postdoctoral Fellow (110243/Z/15/Z).

## References

1. Poolman JT, Wacker M. Extraintestinal pathogenic *Escherichia coli*, a common human pathogen: challenges for vaccine development and progress in the field. J Infect Dis. 2016 213(1):6–13. doi: 10.1093/infdis/jiv429

2. Pitout JDD, DeVinney R. *Escherichia coli* ST131: a multidrug-resistant clone primed for global domination. F1000Research 2017 doi: 10.12688/f1000research.10609.1

3. Goswami C, Fox S, Holden M, Connor M, Leanord A, Evans TJ. Genetic analysis of invasive *Escherichia coli* in Scotland reveals determinants of healthcare-associated versus community-acquired infections. Microb Genom. 2018 4(6). doi: 10.1099/mgen.0.000190.

4. Ender PT, Gajanana D, Johnston B, Clabots C, Tamarkin FJ, Johnson JR. Transmission of an extended-spectrum-beta-lactamase-producing *Escherichia coli* (sequence type ST131) strain between a father and daughter resulting in septic shock and Emphysematous pyelonephritis. J Clin Microbiol. 2009 47(11):3780–2. doi: 10.1128/JCM.01361-09.

5. Van der Bij AK, Peirano G, Pitondo-Silva A, Pitout JD. The presence of genes encoding for different virulence factors in clonally related *Escherichia coli* that produce CTX-Ms. Diagn Microbiol Infect Dis. 2012 72(4):297–302. doi: 10.1016/j.diagmicrobio.2011.12.011

6. Calhau V, Ribeiro G, Mendonça N, Da Silva GJ. Prevalent combination of virulence and plasmidic-encoded resistance in ST131 *Escherichia coli* strains. Virulence. 2013 4(8):726–9. doi: 10.4161/viru.26552.

7. Totsika M, Beatson SA, Sarkar S, Phan MD, Petty NK, Bachmann N, Szubert M, Sidjabat HE, Paterson DL, Upton M, Schembri MA. Insights into a multidrug resistant *Escherichia coli* pathogen of the globally disseminated ST131 lineage: genome analysis and virulence mechanisms. PLoS One. 2011 6(10):e26578. doi: 10.1371/journal.pone.0026578.

8. Ben Zakour NL, Alsheikh-Hussain AS, Ashcroft MM, Khanh Nhu NT, Roberts LW, Stanton-Cook M, Schembri MA, Beatson SA. Sequential acquisition of virulence and fluoroquinolone resistance has shaped the evolution of *Escherichia coli* ST131. MBio. 2016 7(2):e00347–16. doi: 10.1128/mBio.00347-16.

9. Forde BM, Phan MD, Gawthorne JA, Ashcroft MM, Stanton-Cook M, Sarkar S, Peters KM, Chan KG, Chong TM, Yin WF, Upton M, Schembri MA, Beatson SA. Lineage-specific methyltransferases define the methylome of the globally disseminated *Escherichia coli* ST131 clone. MBio. 2015 6(6):e01602–15. doi: 10.1128/mBio.01602-15.

10. Johnson JR, Johnston B, Clabots C, Kuskowski MA, Castanheira M. *Escherichia coli* sequence type ST131 as the major cause of serious multidrug-resistant *E. coli* infections in the United States. Clin Infect Dis. 2010 51(3):286–94. doi: 10.1086/653932.

11. Juhas M, van der Meer JR, Gaillard M, Harding RM, Hood DW, Crook DW. Genomic islands: tools of bacterial horizontal gene transfer and evolution. FEMS Microbiol Rev. 2009 33(2):376–93. doi: 10.1111/j.1574-6976.2008.00136.x

12. Frost LS, Leplae R, Summers AO, Toussaint A. Mobile genetic elements: the agents of open source evolution. Nat Rev Microbiol. 2005 3(9):722–32.

13. Hinnebusch J, Tilly K. Linear plasmids and chromosomes in bacteria. Mol Microbiol. 1993 10(5):917–22.

14. Woodford N, Carattoli A, Karisik E, Underwood A, Ellington MJ, Livermore DM. Complete nucleotide sequences of plasmids pEK204, pEK499, and pEK516, encoding CTX-M enzymes in three major *Escherichia coli* lineages from the United Kingdom, all belonging to the international O25:H4-ST131 clone. Antimicrob Agents Chemother. 2009 53(10):4472–82. doi: 10.1128/AAC.00688-09.

15. Nicolas-Chanoine MH, Bertrand X, Madec JY. *Escherichia coli* ST131, an intriguing clonal group. Clin Microbiol Rev. 2014 27(3):543–74. doi:10.1128/CMR.00125-13.

16. Phan MD, Forde BM, Peters KM, Sarkar S, Hancock S, Stanton-Cook M, Ben Zakour NL, Upton M, Beatson SA, Schembri MA. Molecular characterization of a multidrug resistance IncF plasmid from the globally disseminated *Escherichia coli* ST131 clone. PLoS One. 2015 10(4):e0122369. doi: 10.1371/journal.pone.0122369.

17. Harrison E, Brockhurst MA. Plasmid-mediated horizontal gene transfer is a coevolutionary process. Trends Microbiol. 2012 20(6):262–7. doi: 10.1016/j.tim.2012.04.003

18. MacLean RC, San Millan A. Microbial Evolution: Towards Resolving the Plasmid Paradox. Curr Biol. 2015 25(17):R764–7. doi: 10.1016/j.cub.2015.07.006

19. Shintani M, Sanchez ZK, Kimbara K. Genomics of microbial plasmids: classification and identification based on replication and transfer systems and host taxonomy. Front Microbiol 2015;6.

20. McNally A, Oren Y, Kelly D, Pascoe B, Dunn S, Sreecharan T, Vehkala M, Välimäki N, Prentice MB, Ashour A, Avram O, Pupko T, Dobrindt U, Literak I, Guenther S, Schaufler K, Wieler LH, Zhiyong Z, Sheppard SK, McInerney JO, Corander J. Combined Analysis of Variation in Core, Accessory and Regulatory Genome Regions Provides a Super-Resolution View into the Evolution of Bacterial Populations. PLoS Genet. 2016 12(9):e1006280. doi: 10.1371/journal.pgen.1006280.

21. Wick RR, Judd LM, Gorrie CL, Holt KE. Completing bacterial genome assemblies with multiplex MinION sequencing. Microb Genom. 2017a 3(10):e000132. doi: 10.1099/mgen.0.000132.

22. Arredondo-Alonso S, Rogers MRC, Braat JC, Verschuuren TD, Top J, Corander J, Willems RJL, Schürch AC. mlplasmids: a user-friendly tool to predict plasmid- and chromosome-derived sequences for single species. Microb Genom. 2018 4(11). doi: 10.1099/mgen.0.000224.

23. Judge K, Hunt M, Reuter S, Tracey A, Quail MA, Parkhill J, Peacock SJ. Comparison of bacterial genome assembly software for MinION data and their applicability to medical microbiology. Microb Genom. 2016 2(9):e000085. doi: 10.1099/mgen.0.000085

24. Leggett RM, Clark MD. A world of opportunities with nanopore sequencing. J Exp Bot. 2017 68(20):5419–5429. doi: 10.1093/jxb/erx289.

25. Roer L, Overballe-Petersen S, Hansen F, Johannesen TB, Stegger M, Bortolaia V, Leekitcharoenphon P, Korsgaard HB, Seyfarth AM, Mossong J, Wattiau P, Boland C, Hansen DS, Hasman H, Hammerum AM, Hendriksen RS. ST131 fimH22 *Escherichia coli* isolate with a blaCMY-2/IncI1/ST12 plasmid obtained from a patient with bloodstream infection: highly similar to *E. coli* isolates of broiler origin. JAntimicrob Chemother. 2018 doi: 10.1093/jac/dky484.

26. Goldstein S, Beka L, Graf J, Klassen J. Evaluation of strategies for the assembly of diverse bacterial genomes using MinION long-read sequencing. 2018 Biorxiv doi: https://doi.org/10.1101/362673

27. Ludden C, Reuter S, Judge K, Gouliouris T, Blane B, Coll F, Naydenova P, Hunt M, Tracey A, Hopkins KL, Brown NM, Woodford N, Parkhill J, Peacock SJ. Sharing of carbapenemase-encoding plasmids between *Enterobacteriaceae* in UK sewage uncovered by MinION sequencing. Microb Genom. 2017 3(7):e000114. doi: 10.1099/mgen.0.000114.

28. Lanfear R, Schalamun M, Kainer D, Wang W, Schwessinger B. MinIONQC: fast and simple quality control for MinION sequencing data. Bioinformatics. 2018 doi: 10.1093/bioinformatics/bty654

29. Wick RR, Judd LM, Gorrie CL, Holt KE. Unicycler: Resolving bacterial genome assemblies from short and long sequencing reads. PLoS Comput Biol. 2017b 13(6):e1005595. doi: 10.1371/journal.pcbi.1005595.

30. Ewels P, Magnusson M, Lundin S, Käller M. MultiQC: summarize analysis results for multiple tools and samples in a single report. Bioinformatics. 2016 32(19):3047–8. doi: 10.1093/bioinformatics/btw354.

31. George S, Pankhurst L, Hubbard A, Votintseva A, Stoesser N et al. Resolving plasmid structures in *Enterobacteriaceae* using the MinION nanopore sequencer: assessment of MinION and MinION/Illumina hybrid data assembly approaches. Microb Genom 2017:1–8.

32. Gurevich A, Saveliev V, Vyahhi N, Tesler G. QUAST: quality assessment tool for genome assemblies. Bioinformatics. 2013 29(8):1072–5. doi: 10.1093/bioinformatics/btt086.

33. Wick RR, Schultz MB, Zobel J, Holt KE. Bandage: interactive visualization of de novo genome assemblies. Bioinformatics. 2015 31(20):3350–2. doi: 10.1093/bioinformatics/btv383.

34. Waterhouse RM, Seppey M, Simão FA, Manni M, Ioannidis P, Klioutchnikov G, Kriventseva EV, Zdobnov EM. BUSCO applications from quality assessments to gene prediction and phylogenomics. Mol Biol Evol. 2017 doi: 10.1093/molbev/msx319.

35. Seemann T. Prokka: rapid prokaryotic genome annotation. Bioinformatics. 2014 30(14):2068–9. doi: 10.1093/bioinformatics/btu153

36. Partridge SR, Tsafnat G. Automated annotation of mobile antibiotic resistance in gram-negative bacteria: The Multiple Antibiotic Resistance Annotator (MARA) and database. J Antimicrob Chemother. 2018 73(4):883–890. doi: 10.1093/jac/dkx513.

37. Carattoli A, Zankari E, García-Fernández A, Voldby Larsen M, Lund O, Villa L, Møller Aarestrup F, Hasman H. In silico detection and typing of plasmids using PlasmidFinder and plasmid multilocus sequence typing. Antimicrob Agents Chemother. 2014 58(7):3895–903. doi: 10.1128/AAC.02412-14.

38. Brooks L, Kaze M, Sistrom M. A Curated, Comprehensive Database of Plasmid Sequences. Microbiol Resour Announc. 2019 8(1). pii: e01325–18. doi: 10.1128/MRA.01325-18

39. Sullivan MJ, Petty NK, Beatson SA. Easyfig: a genome comparison visualizer. Bioinformatics. 2011 27(7):1009–10. doi: 10.1093/bioinformatics/btr039

40. Matsumura Y, Pitout JDD, Peirano G, DeVinney R, Noguchi T, Yamamoto M, Gomi R, Matsuda T, Nakano S, Nagao M, Tanaka M, Ichiyama S. Rapid identification of different *Escherichia coli* sequence type 131 clades. Antimicrob Agents Chemother. 2017; 61(8). pii: e00179–17. doi: 10.1128/AAC.00179-17.

41. Chen S, Zhou Y, Chen Y, Gu J. Fastp: an ultra-fast all-in-one FASTQ preprocessor. Bioinformatics. 2018 34(17):i884–i890.

42. Page AJ, Cummins CA, Hunt M, Wong VK, Reuter S, Holden MT, Fookes M, Falush D, Keane JA, Parkhill J. Roary: rapid large-scale prokaryote pan genome analysis. Bioinformatics. 2015 31(22):3691–3. doi: 10.1093/bioinformatics/btv421

43. Katoh K, Standley DM. MAFFT multiple sequence alignment software version 7: improvements in performance and usability. Mol Biol Evol. 2013 30(4):772–80. doi: 10.1093/molbev/mst010

44. Stamatakis A. RAxML version 8: a tool for phylogenetic analysis and post-analysis of large phylogenies. Bioinformatics. 2014 30(9):1312–3. doi: 10.1093/bioinformatics/btu033

45. Partridge SR, Kwong SM, Firth N, Jensen SO. Mobile genetic elements associated with antimicrobial resistance. Clinical Microbiology Reviews 31(4):e00088–17 2018 doi: 10.1128/CMR.00088-17

46. Lartigue MF, Poirel L, Aubert D, Nordmann P. In vitro analysis of IS*Ecp1B*-mediated mobilization of naturally occurring beta-lactamase gene blaCTX-M of *Kluyvera ascorbata*. Antimicrob Agents Chemother. 2006 50(4):1282–6.

47. Barlow M, Reik RA, Jacobs SD, Medina M, Meyer MP, McGowan JE Jr, Tenover FC. High rate of mobilization for blaCTX-Ms. Emerg Infect Dis. 2008 14(3):423–8. doi: 10.3201/eid1403.070405.

48. Petty NK, Ben Zakour NL, Stanton-Cook M, Skippington E, Totsika M, Forde BM, Phan MD, Gomes Moriel D, Peters KM, Davies M, Rogers BA, Dougan G, Rodriguez-Baño J, Pascual A, Pitout JD, Upton M, Paterson DL, Walsh TR, Schembri MA, Beatson SA. Global dissemination of a multidrug resistant *Escherichia coli* clone. Proc Natl Acad Sci U S A. 2014 111(15):5694–9. doi: 10.1073/pnas.1322678111

49. Lemon JK, Khil PP, Frank KM, Dekker JP. Rapid Nanopore Sequencing of Plasmids and Resistance Gene Detection in Clinical Isolates. J Clin Microbiol. 2017 55(12):3530–3543. doi: 10.1128/JCM.01069-17.

50. Stoesser N, Batty EM, Eyre DW, Morgan M, Wyllie DH, Del Ojo Elias C, Johnson JR, Walker AS, Peto TEA, Crook DW. Predicting antimicrobial susceptibilities for *Escherichia coli* and *Klebsiella pneumoniae* isolates using whole genomic sequence data. J Antimicrob Chemother 2013 68:2234–2244.

51. Branger C, Ledda A, Billard-Pomares T, Doublet B, Fouteau S, Barbe V, Roche D, Cruveiller S, Médigue C, Castellanos M, Decré D, Drieux-Rouze L, Clermont O, Glodt J, Tenaillon O, Cloeckaert A, Arlet G, Denamur E. Extended-spectrum β-lactamase-encoding genes are spreading on a wide range of *Escherichia coli* plasmids existing prior to the use of third-generation cephalosporins. Microb Genom. 2018 4(9). doi: 10.1099/mgen.0.000203.

52. Moradigaravand D, Palm M, Farewell A, Mustonen V, Warringer J, Parts L. Prediction of antibiotic resistance in *Escherichia coli* from large-scale pan-genome data. 2018 PLoS Computational Biology doi: https://doi.org/10.1371/journal.pcbi.1006258.

53. Greig DR, Dallman TJ, Hopkins KL, Jenkins C. MinION nanopore sequencing identifies the position and structure of bacterial antibiotic resistance determinants in a multidrug-resistant strain of enteroaggregative *Escherichia coli*. Microb Genom. 2018 4(10). doi: 10.1099/mgen.0.000213.

54. Wang Y, Yang Q, Wang Z. The evolution of nanopore sequencing. Front Genet 2014 5:449.

55. Tamma PD, Y Fan, Bergman Y, Pertea G, Kazmi A, Lewis S, Carroll KC, Schatz MC, Timp W, Simner P. Rapid optimization of antibiotic therapy for multidrug-resistant gram-negative infections using Nanopore whole genome sequencing. 2018 Available at SSRN: https://ssrn.com/abstract=3219539.

56. Tyson GH, McDermott PF, Li C, Chen Y, Tadesse DA, Mukherjee S, Bodeis-Jones S, Kabera C, Gaines SA, Loneragan GH, Edrington TS, Torrence M, Harhay DM, Zhao S. WGS accurately predicts antimicrobial resistance in *Escherichia coli*. J Antimicrob Chemother 2015 70:2763–2769.

57. Su M, Satola SW, Read TD. Genome-based prediction of bacterial antibiotic resistance. J Clin Microbiol. 2018 doi: 10.1128/JCM.01405-18.

58. Lutgring JD, Zhu W, de Man TJB, Avillan JJ, Anderson KF, Lonsway DR, Rowe LA, Batra D, Rasheed JK, Limbago BM. Phenotypic and Genotypic Characterization of *Enterobacteriaceae* Producing Oxacillinase-48-Like Carbapenemases, United States. Emerg Infect Dis. 2018 24(4):700–709. doi: 10.3201/eid2404.171377.

