## Supplementary_Data for "Complete assembly of *Escherichia coli* ST131 genomes using long reads demonstrates antibiotic resistance gene variation within diverse plasmid and chromosomal contexts"

**Table of contents**


| | **Page** |
| --- | --- |
| Supplementary Results | 1 |
| Supplementary Figure 1 | 2 |
| Supplementary Figure 2 | 3-5 |
| Supplementary Figure 3 | 6 |
| Supplementary Figure 4 | 7 |
| Supplementary Figure 5 | 8 |
| Supplementary Figure 6 | 9 |
| Supplementary Table 1 | 10 |
| Supplementary Table 2 | 10 |
| Supplementary Table 3 | 11 |
| Supplementary Table 4 | 12 |


### Supplementary Results

#### Oxford Nanopore long read quality control and long read genome assembly

Half of the reads were produced within 14 hours of sequencing, with the remainder produced over the subsequent 34 hours (Supplementary Figure 1d). A median read length of 5.5 Kb for reads Q (quality) score $\geq$ 7 was achieved within one hour of sequencing (Supplementary Figure 1e), and the median Q score declined slightly as the run proceeded (Supplementary Figure 1f). An average of 30-fold theoretical coverage from 954 Mbases with Q $\geq$ 7 was exceeded in this GridION run within three hours.

We compared short read-only, long read-only and hybrid assembly outputs from Unicycler v.4.6 using the long Oxford Nanopore reads and short Illumina reads to identify the most contiguous assemblies per sample across all three Unicycler modes (conservative, normal and bold).

For five samples, the long read assemblies produced 2-7 contigs (with a median of three) with nearly identical results across modes, whereas the short read assemblies resulted in 76-230 contigs (a median of 124), and the hybrid assemblies also had more contigs (6-191 with a median of 44). For VREC0739 and VREC1428, the short read libraries resulted in over-bridging of contigs making it harder to classify contigs as chromosomal or plasmid-associated, perhaps because long reads already provided sufficient genome coverage and the assembler inserted the contigs produced by short reads at short homologous repetitive regions.

#### VREC1013 assembly assessment and improvement

For VREC1013, the hybrid assembly improved the long read assembly such that the final optimised version had three rather than 22 contigs and a smaller length (5.36 Mb, Supplementary Table 3), after manual sequence alignment eliminated seven false-positive short contigs. Five contigs had depths of coverage <8% of the chromosomal median and may were the result of contig overbridging during assembly. Pairwise alignment of these five contigs with BLAST against the assembly showed that they had near-perfect matches (E-value < E-10) with other contigs, showing that they were effectively duplicate contigs, and thus few reads mapped to them. In contrast, the other four valid contigs acted positive controls and showed high homology to their own contigs only. As a result, duplicate contigs were removed from the VREC1013 hybrid assembly used for subsequent analyses.

Supplementary Figure 1. Overview of genome assembly using Oxford Nanopore reads to recover plasmids with antibiotic resistance genes and mobile genetic elements (MGEs). Oxford Nanopore fast5 sequences were basecalled and converted to fastq format using Albacore v.2.0 and Guppy v.0.5.1. Forward, reverse and middle adapters were removed using Porechop v.0.2.4. The genomes were assembled using Unicycler v.4.6 (optionally including Illumina short reads for comparison). The probability that the resulting contigs were chromosomal or plasmid-associated was measured using mlplasmids. Contigs were annotated using the Comprehensive Antibiotic Resistance Database (CARD) and Multiple antibiotic Resistance Annotator (MARA) to resolve precise plasmid structure, *bla*$_{CTX-M}$ gene alleles, copy numbers and their adjacent regions.

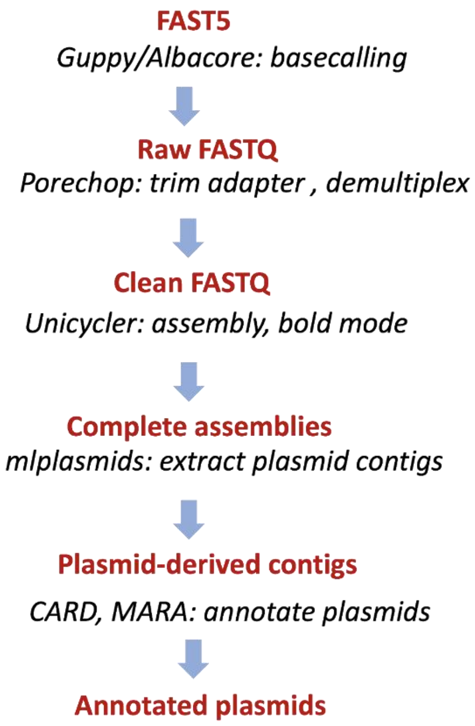

Supplementary Figure 2a-g. Summary plots of the GridION X5 sequencing run for all (blue) and filtered (green) nanopore reads generated using MinIONQC. The graphs in (a) show the read count (y-axis) with the mean and median read length and the number of bases and reads per channel (x-axis), the overall read count (y-axis) vs length (x-axis) in (b) and read count (y-axis) vs the mean Q score (x-axis) in (c). Plots were also drawn to present the total amount of bases called (x-axis; d), the mean read length (x-axis; e) and the mean Q score (x-axis; f) per hour (in their y-axes); the total amount of bases (y-axis) contained in a minimum read length (x-axis) is shown in (g).

(a)

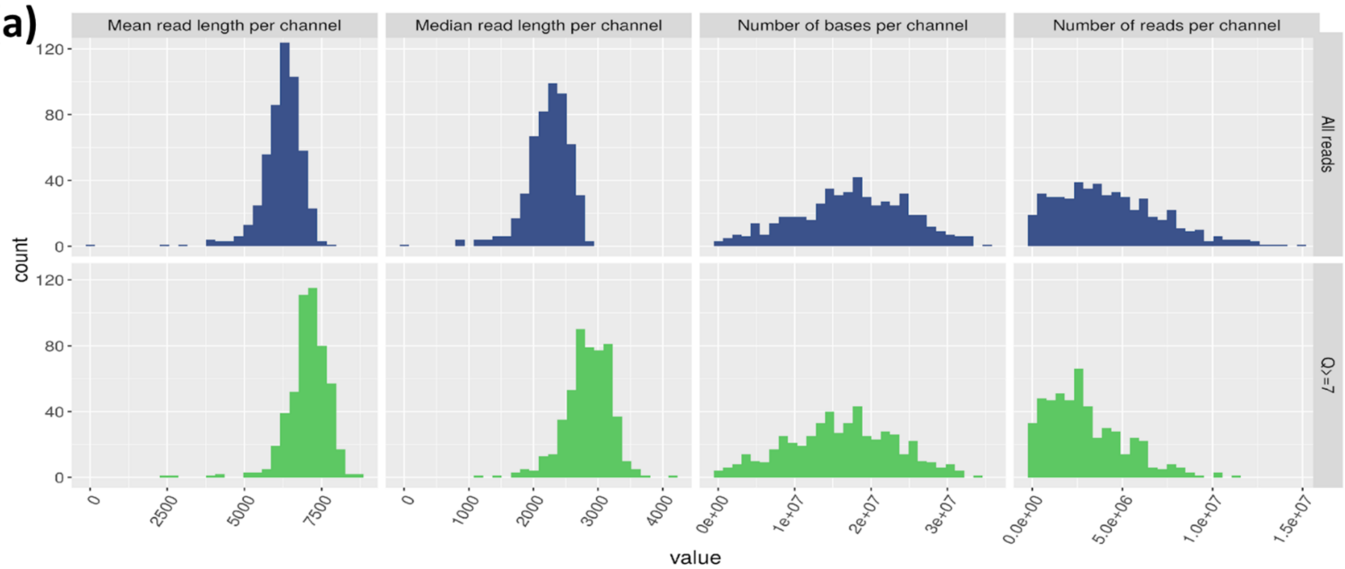


(b)

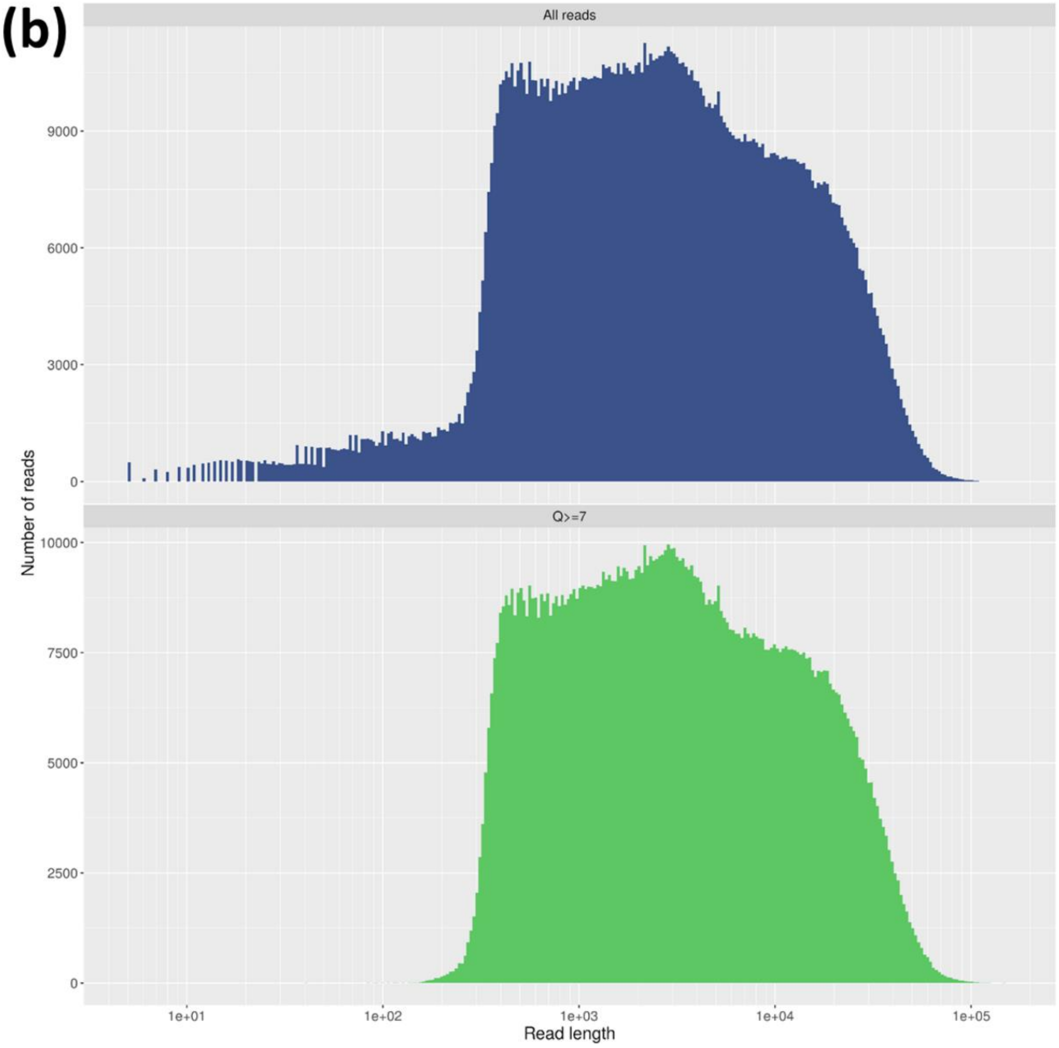

(c)

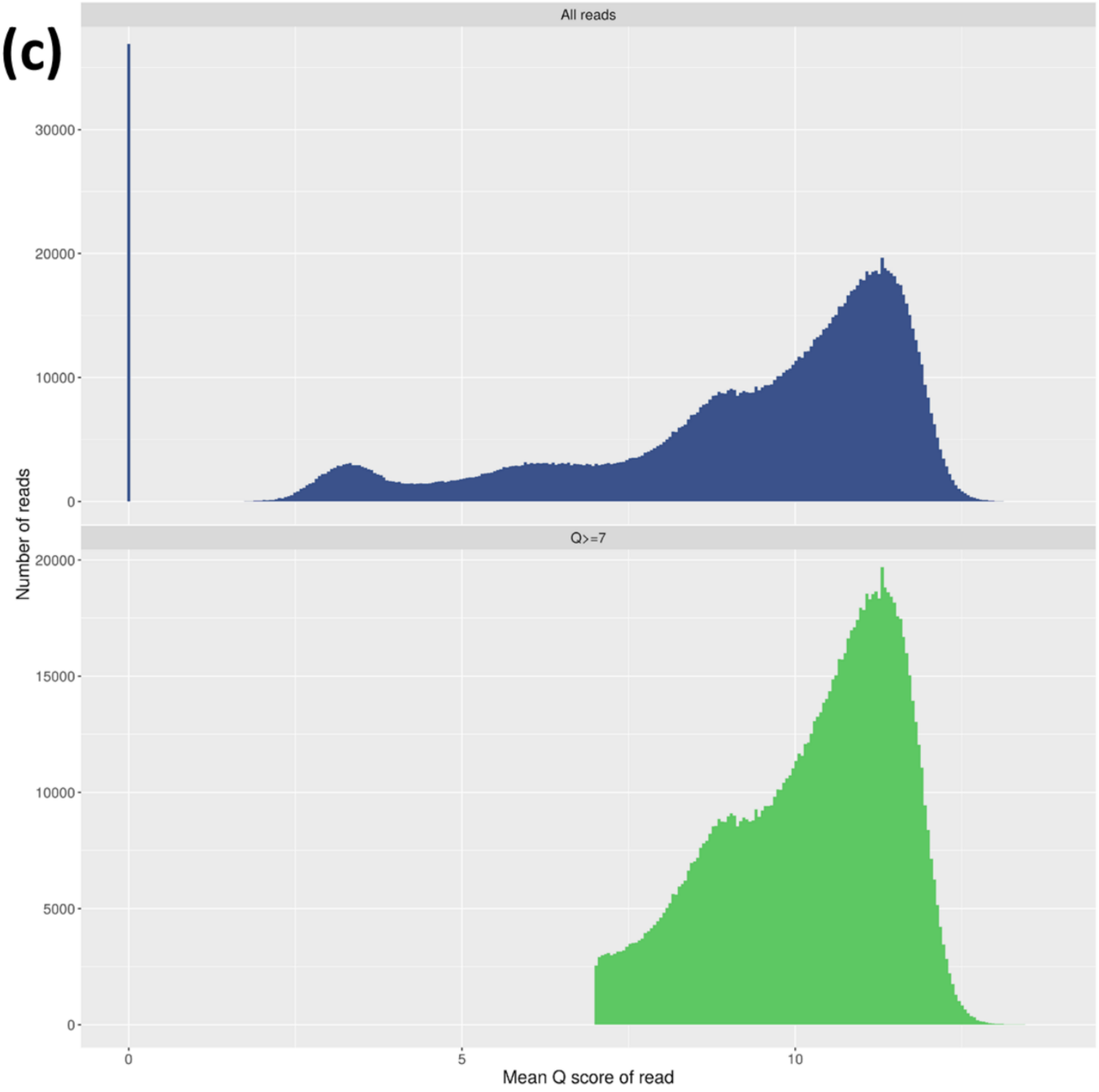
All reads
Q>=7
Number of reads
30000
20000
10000
0
15000
5000
0
5
10
Mean Q score of read

(d)

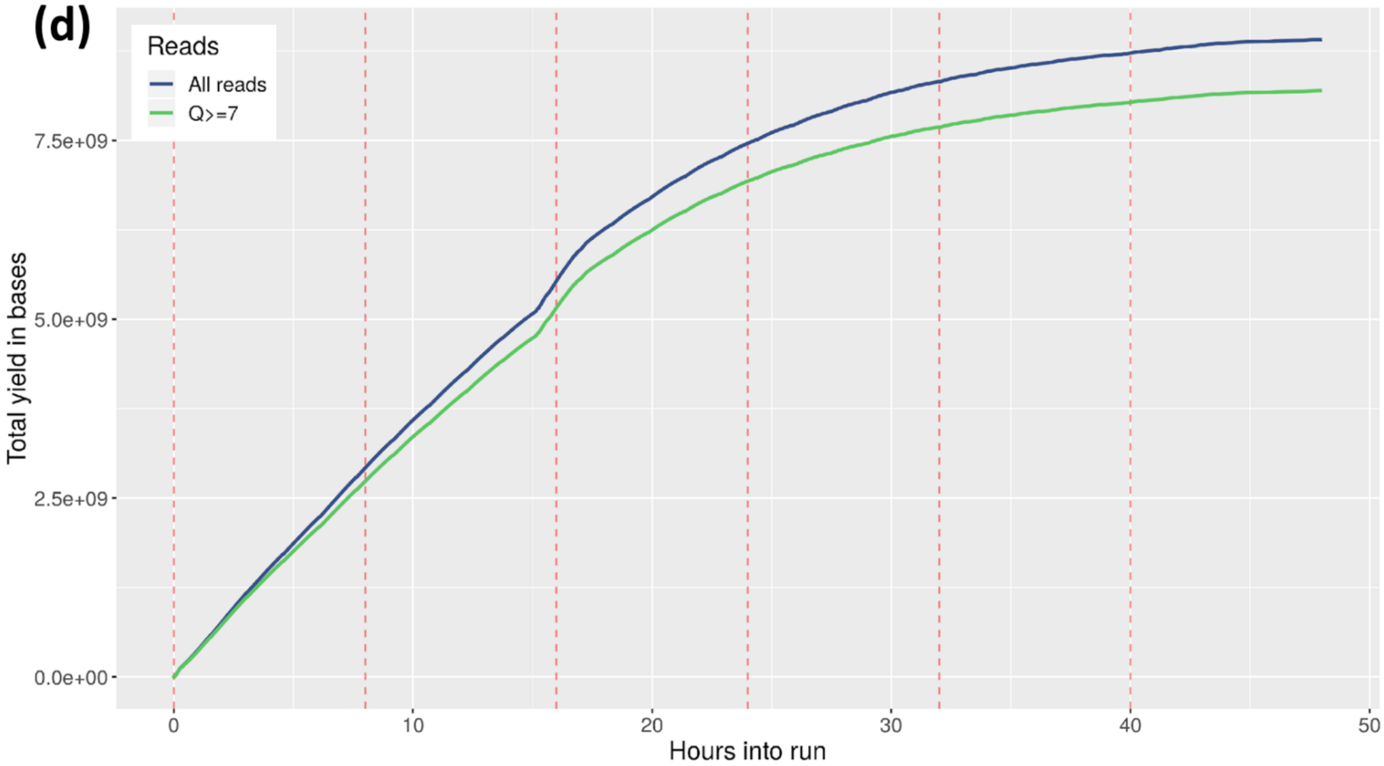
Reads
All reads
Q>=7
7.5e+09
5.0e+09
2.5e+09
0.0e+00
Total yield in bases
0
10
20
30
40
50
Hours into run

(e)

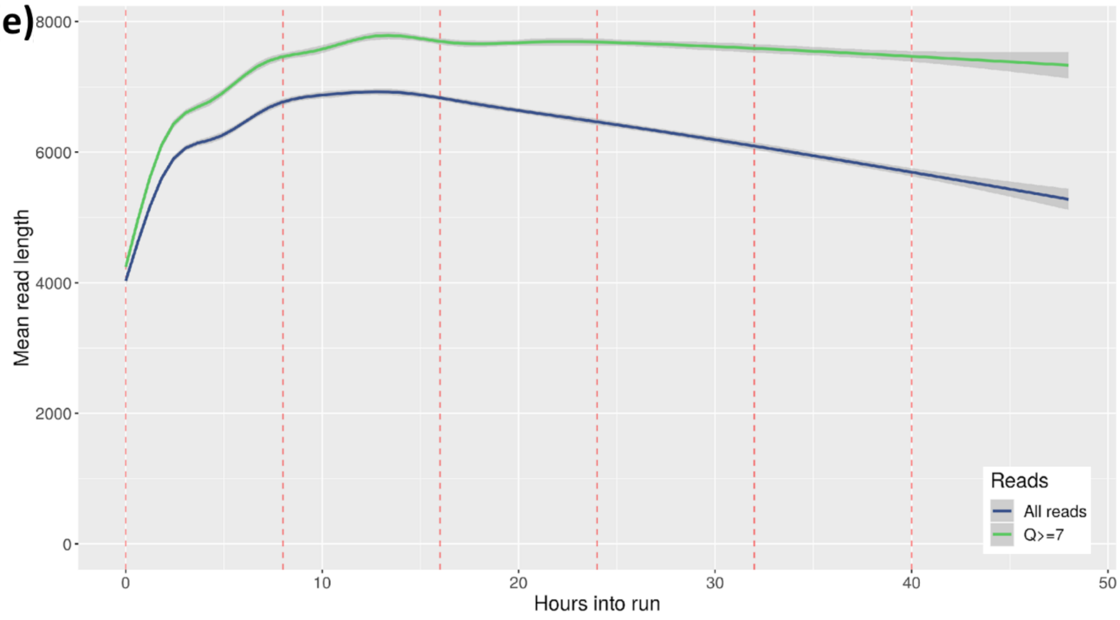
8000
6000
4000
2000
0
Mean read length
0
10
20
30
40
50
Hours into run
Reads
All reads
Q>=7

(f)

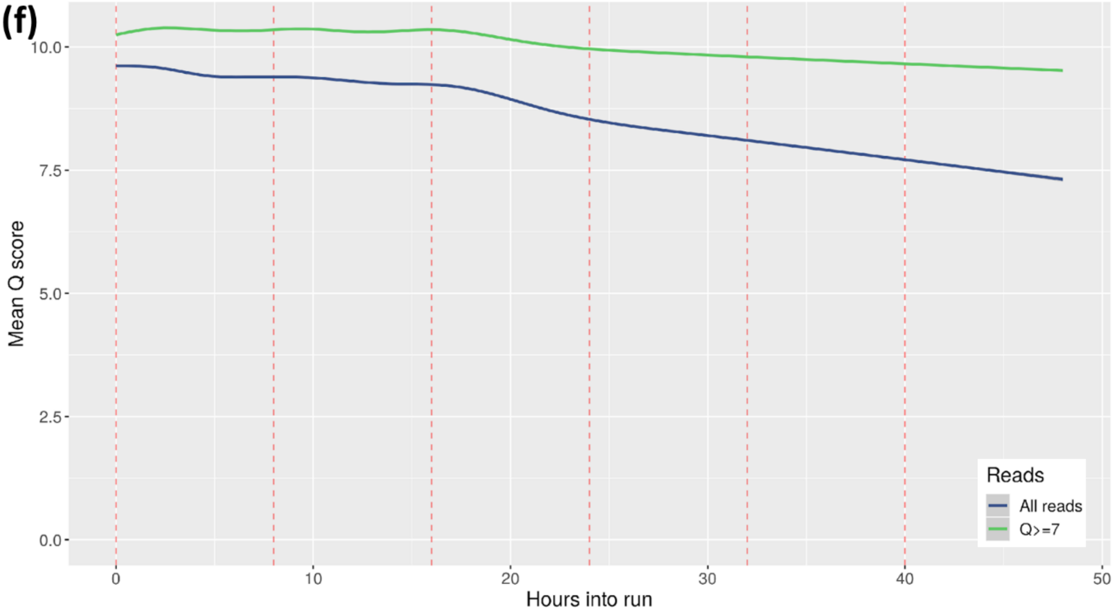
10.0
7.5
5.0
2.5
0.0
Mean Q score
0
10
20
30
40
50
Hours into run
Reads
All reads
Q>=7

(g)

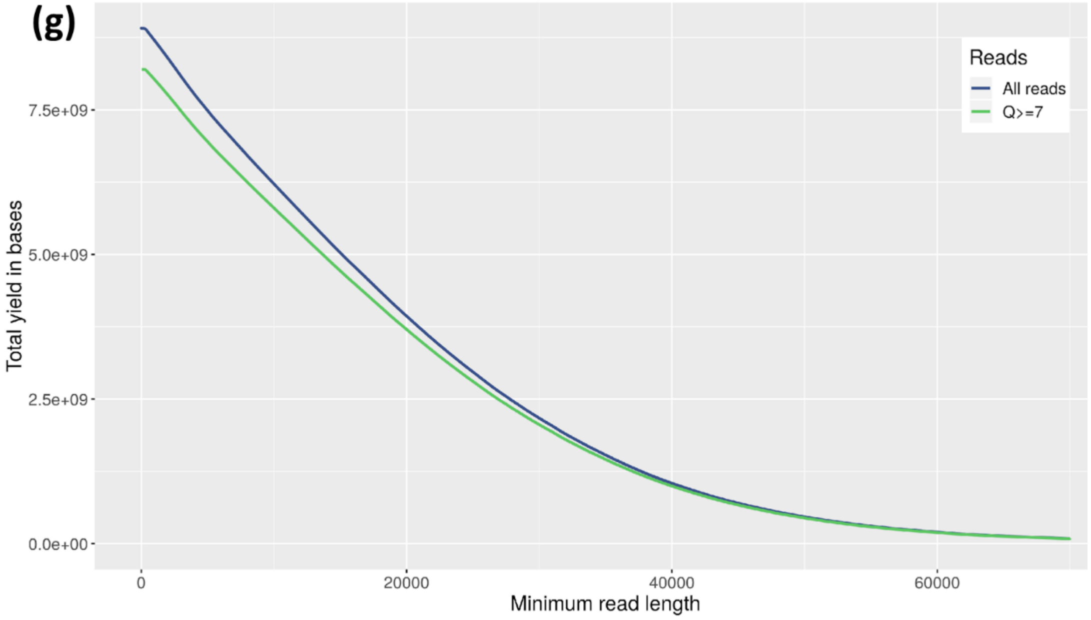
7.5e+09
5.0e+09
2.5e+09
0.0e+00
Total yield in bases
0
20000
40000
60000
Minimum read length
Reads
All reads
Q>=7

Supplementary Figure 3. Summary of the GridION X5 sequencing run output showing the read length on a log10 scale (x-axis) versus the mean Q score of each read (y-axis) where points are coloured by events per base. The horizontal red line shows reads with lengths > 10 Kb and the vertical red line read with Q scores > 10. Together, this area shows the large number of long high-quality reads generated in this study. This plot emphasises that a high proportion of the bases were accurately called: these were subsequently used for downstream analysis.

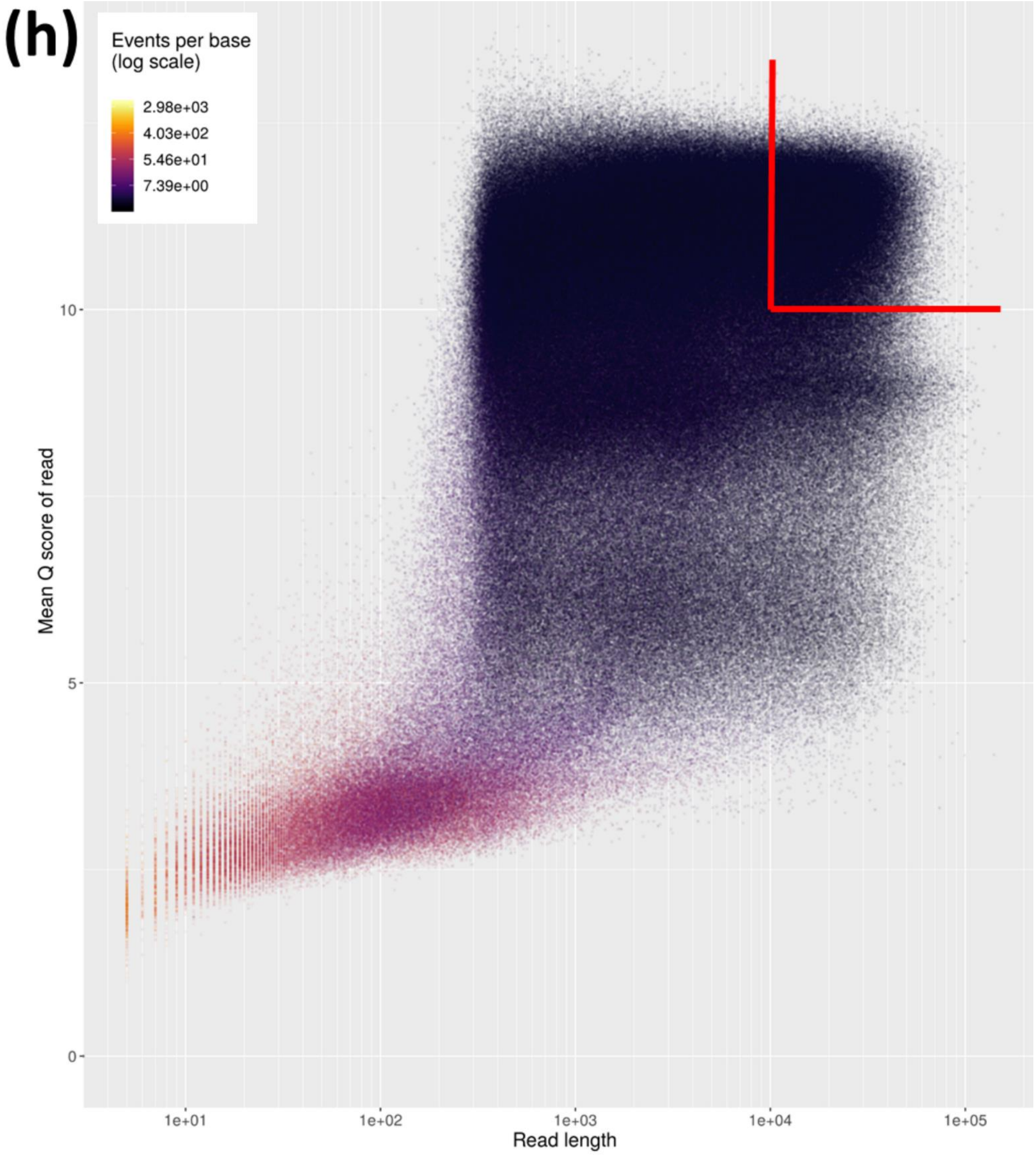

Supplementary Figure 4. The assembly graphs of six *E. coli* ST131 genomes showed many connected edges for those created from short Illumina HiSeq reads only (left) but near-complete assemblies for those made with long Oxford Nanopore read-only (centre) and the hybrid assemblies of most of the strains (right). The assemblies were generated with Unicycler v.4.6 and were visualised using Bandage. Circularized contigs indicated complete assemblies.

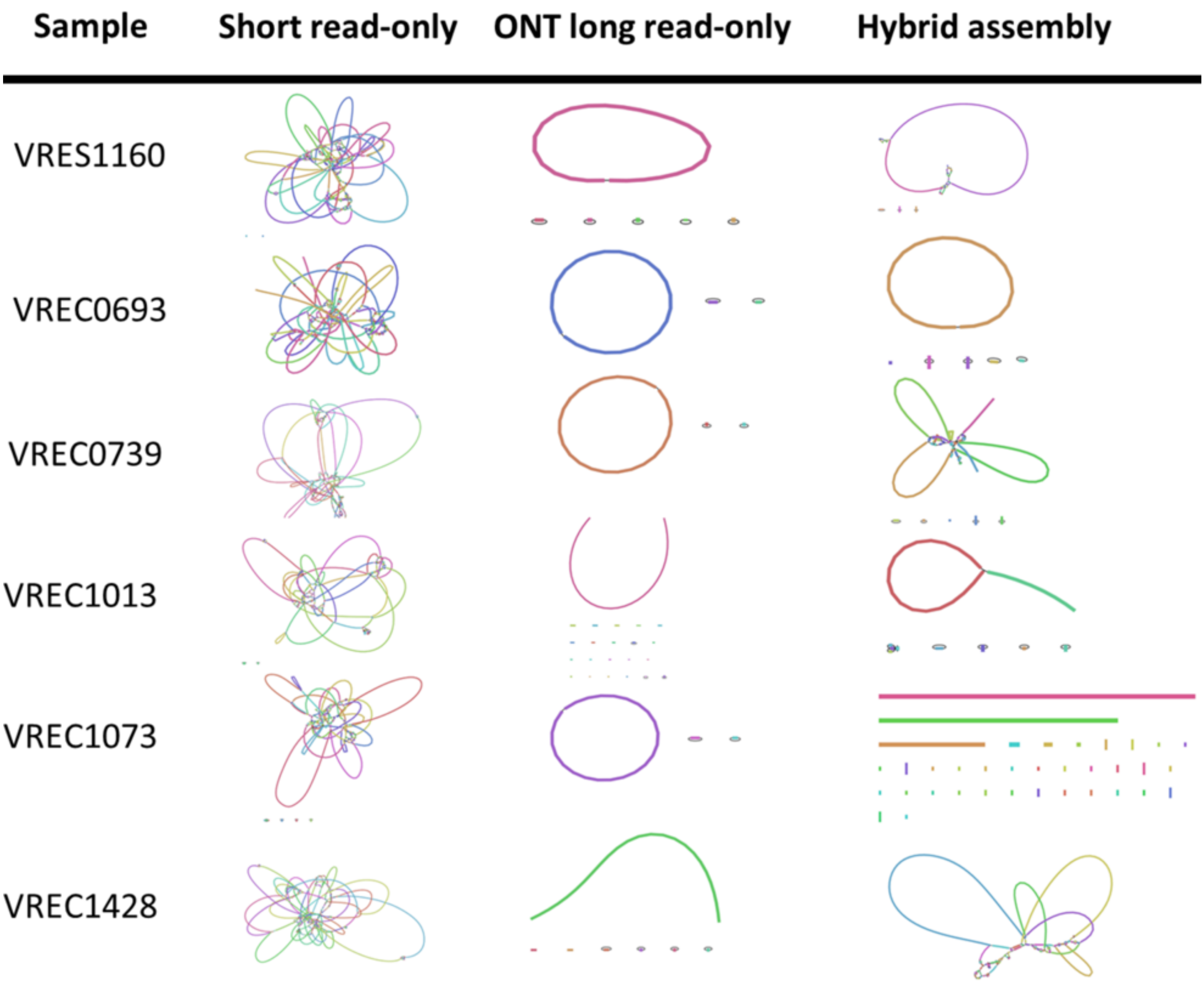

Supplementary Figure 5. The contigs from the most optimal assembly mode of Unicycler v.4.6 of five out of six *E. coli* ST131 samples were identified as chromosomal or plasmid-derived using mlplasmids. These were annotated with *bla*$_{CTX-M}$ genes and their genetic flanking context using Galileo™ AMR based on the Multiple Antibiotic Resistance Annotator (MARA) and database [35]; all *bla*$_{CTX-M}$ variants are labelled accordingly and encircled in red (*bla*$_{CTX-M-15}$), purple (*bla*$_{CTX-M-14}$) or green (*bla*$_{CTX-M-27}$). The definition of the other elements are listed at https://galileoamr.arcbio.com/mara/feature/list. The long VREC0693 chromosome is split into two parts so that the gene annotation is visible.

VRES1160 (subclade C2, 61,934 bp *bla*$_{CTX-M-15}$+ plasmid)

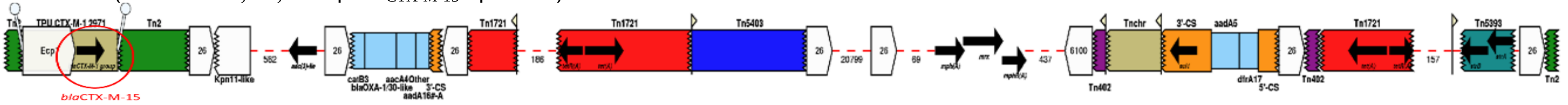


VREC1013 (subclade C2, 89,945 bp *bla*$_{CTX-M-15}$+ plasmid)

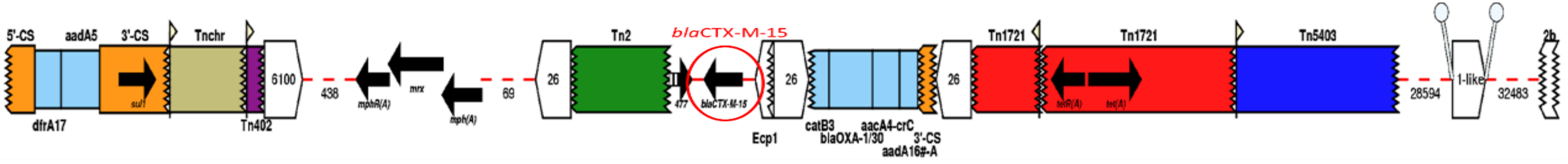


VREC1428 (subclade C1, 92,750 bp *bla*$_{CTX-M-27}$+ plasmid)

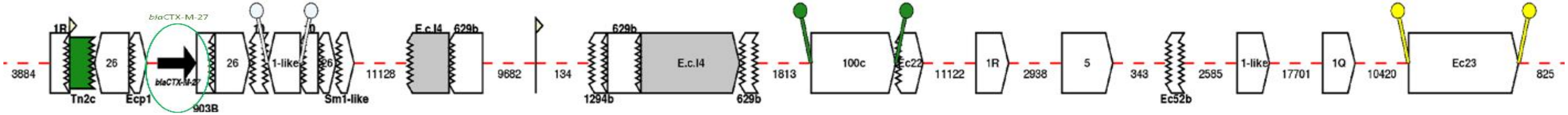


VREC0693 (subclade C2, 5,039,909 bp *bla*$_{CTX-M-15}$+ chromosome with 3 distinct *bla*$_{CTX-M-15}$ genes in red - single chromosome is split below for visualisation)

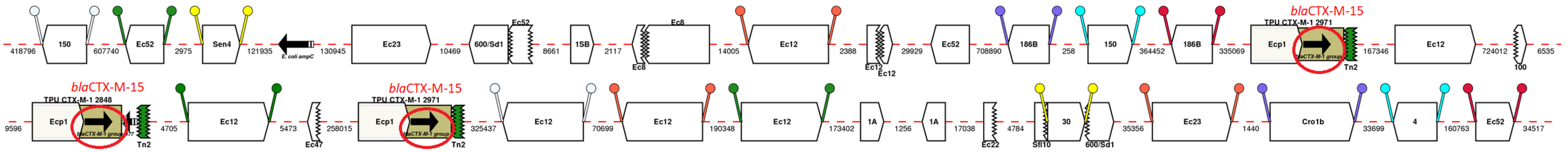


VREC1073 (subclade C2, 96,056 bp *bla*$_{CTX-M-14}$+)

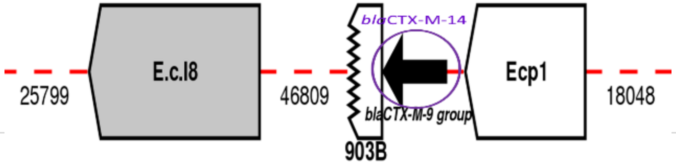

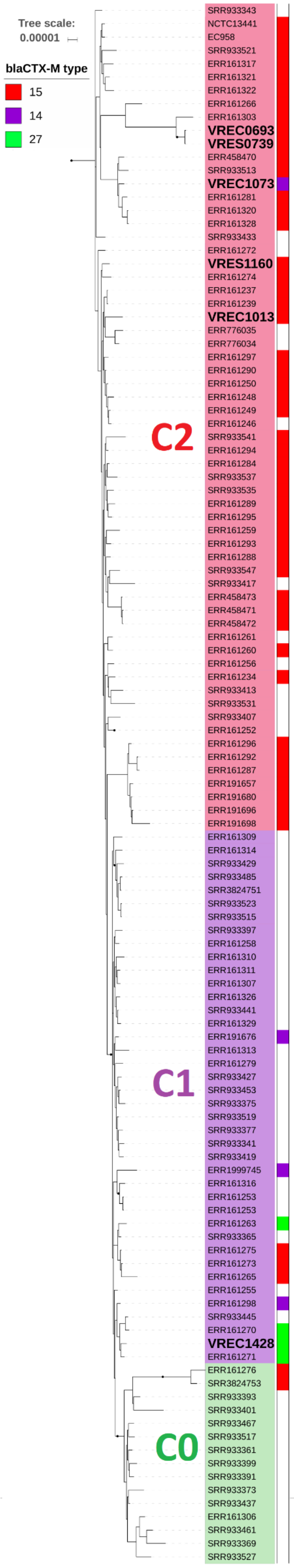


Supplementary Figure 6. Phylogram of the six ST131 genomes showed that all except VREC1428 were in ST131 subclade C2 (red: VRES1160, VREC1073, VRES0739, VREC0693 and VREC1013). VREC1428 clustered in subclade C1 (purple). No new isolate was in C0 (green). The phylogram was built with RAxML v.8.2.11 and iTOL v4.3 using 3,603 non-recombinant SNPs from Gubbins v.2.3.4 where branch support was performed by 100 bootstrap replicates, and the scale bar indicates the number of substitutions per site. Clade classification was based on phylogenetic analysis by [8] by including the reference NCTC13441, n=63 isolates from [8] and n=56 from [42] with associated classification and *bla*$_{CTX-M}$ allele data. The right-hand part shows *bla*$_{CTX-M-15}$ (red), *bla*$_{CTX-M-14}$ (purple) and *bla*$_{CTX-M-27}$ alleles (green). The six isolates' names are in large bold text. This mid-pointed rooted phylogeny included reference genome isolates EC958 and NCTC13441 (both in C2) and a clade B isolate as an outgroup (Figure 3). The C2 isolates were mainly *bla*$_{CTX-M-15}$-positive (48 out of 62, including VRES1160, VRES0739, VREC0693 and VREC1013), bar 13 that were *bla*$_{CTX-M-}$-negative and one that was *bla*$_{CTX-M-14}$-positive (VREC1073). The C0 isolates were mainly *bla*$_{CTX-M-15}$-negative (13 out of 15), as were the C1 (30 out of 40) isolates except for four that were *bla*$_{CTX-M-27}$-positive, three that were *bla*$_{CTX-M-15}$-positive and three that were *bla*$_{CTX-M-14}$-positive.

Supplementary Table 1. Sample collection source, sampling date and sequence read accession numbers.

| Strain | Source | Sampling date | Accession numbers | |
|---|---|---|---|---|
| | | | Short reads | Long reads |
| VRES1160 | Faeces | 26/08/2015 | ERR1878359 | https://ndownloader.figshare.com/files/14039495 |
| VREC0693 | Faeces | 03/06/2015 | ERR2137889 | https://ndownloader.figshare.com/files/14039639 |
| VRES0739 | Faeces | 05/06/2015 | ERR1878196 | https://ndownloader.figshare.com/files/14039354 |
| VREC1013 | Faeces | 19/08/2015 | ERR2138591 | https://ndownloader.figshare.com/files/14039333 |
| VREC1073 | Blood | 26/08/2015 | ERR2138200 | https://ndownloader.figshare.com/files/14039345 |
| VREC1428 | Faeces | 22/10/2015 | ERR2138475 | https://ndownloader.figshare.com/files/14039351 |

Supplementary Table 2. Contigs were classified as chromosomal or plasmid-derived using the mlplasmids prediction value. Each contig were aligned against CARD to identify the presence/absence of *bla*$_{CTX\text{-}M}$ alleles and their copy numbers. Plasmid types were identified using PlasmidFinder.

| Strain | Prediction | Prediction value (%) | Contig ID | Length (bp) | *bla*$_{CTX\text{-}M}$ allele | *bla*$_{CTX\text{-}M}$ count | Plasmid type | Median Depth | Normalized Depth |
|---|---|---|---|---|---|---|---|---|---|
| VRES1160 | Chromosome | 98 | 1 | 5,126,679 | | | - | 258 | 1.00 |
| | Chromosome | 70 | 2 | 113,086 | | | - | 213 | 1.00 |
| | Plasmid | 70 | 3 | 61,934 | 15 | 1 | IncFIA | 282 | 1.10 |
| | Plasmid | 85 | 4 | 15,803 | | | ColRNAI | 420 | 1.64 |
| | Plasmid | 81 | 5 | 5,203 | | | ColRNAI | 11 | 0.04 |
| | Plasmid | 83 | 6 | 4,096 | | | Col8282 | 473 | 1.85 |
| VREC0693 | Chromosome | 98 | 1 | 5,039,909 | 15 | 3 | - | 258 | 1.00 |
| | Plasmid | 61 | 2 | 132,042 | | | IncFIB | 213 | 0.83 |
| | Plasmid | 60 | 3 | 88,790 | | | IncB | 282 | 1.09 |
| VRES0739 | Chromosome | 98 | 1 | 4,797,749 | | | - | 171 | 1.00 |
| | Plasmid | 96 | 2 | 5,162 | | | Col156 | 436 | 2.55 |
| | Plasmid | 74 | 3 | 4,001 | | | - | 303 | 1.77 |
| VREC1013 | Chromosome | 97 | 1 | 3,699,451 | | | - | 300 | 1.00 |
| | Chromosome | 97 | 2 | 1,434,037 | | | - | 335 | 1.00 |
| | Plasmid | 84 | 4 | 89,945 | 15 | 1 | IncFII | 1015 | 3.27 |
| VREC1073 | Chromosome | 98 | 1 | 5,286,804 | | | - | 214 | 1.00 |
| | Plasmid | 68 | 2 | 156,298 | | | IncFIA | 172 | 0.80 |
| | Chromosome | 60 | 3 | 96,056 | 14 | 1 | - | 213 | 1 |
| VREC1428 | Chromosome | 98 | 1 | 4,924,536 | | | - | 126 | 1.00 |
| | Chromosome | 97 | 2 | 103,034 | | | - | 57 | 1.00 |
| | Chromosome | 96 | 3 | 101,160 | | | - | 41 | 1.00 |
| | Plasmid | 64 | 4 | 92,750 | 27 | 1 | IncFIA | 85 | 0.67 |
| | Plasmid | 92 | 5 | 5,147 | | | ColRNAI | 168 | 1.33 |
| | Plasmid | 99 | 6 | 5,143 | | | Col156 | 207 | 1.64 |
| | Plasmid | 73 | 7 | 4,649 | | | ColRNAI | 239 | 1.90 |

Supplementary Table 3. Comparison of short read-only, long read-only and hybrid genome assemblies generated using the conservative, normal and bold modes of Unicycler v.04.6. Assemblies were assessed according to their total length, number of contigs produced, N50 (bp), numbers of mismatches per 100 Kb and numbers of indels per 100 Kb.

| Assembly | Mode | Metric | VRES1160 | VREC0693 | VRES0739 | VREC1013 | VREC1073 | VREC1428 |
|---|---|---|---|---|---|---|---|---|
| Short read-only | Conservative | Total length (bp) | 5,142,342 | 5,146,205 | 5,181,497 | 5,208,807 | 4,967,093 | 5,375,468 |
| | | Number of contigs | 168 | 159 | 200 | 148 | 117 | 230 |
| | | N50 (bp) | 124,175 | 132,865 | 138,725 | 134,439 | 157,528 | 135,303 |
| | | #mismatches / 100 Kb | 1.32 | 1.32 | 65.4 | 1.5 | 285.81 | 0.69 |
| | | #indels / 100 Kb | 0.06 | 0.02 | 1.84 | 0.08 | 261.91 | 0.04 |
| | Normal | Total length (bp) | 5,158,728 | 5,171,710 | 5,227,751 | 5,240,888 | 4,989,316 | 5,416,180 |
| | | Number of contigs | 110 | 106 | 123 | 94 | 76 | 148 |
| | | N50 (bp) | 206,138 | 190,908 | 213,071 | 189,184 | 222,158 | 170,443 |
| | | #mismatches / 100 Kb | 4.64 | 0.93 | 69.86 | 4.25 | 284.14 | 2.96 |
| | | #indels / 100 Kb | 0.21 | 0.14 | 2.33 | 0.36 | 262.1 | 0.04 |
| | Bold | Total length (bp) | 5,159,662 | 5,163,846 | 5,207,686 | 5,226,735 | 4,977,746 | 5,411,973 |
| | | Number of contigs | 124 | 120 | 146 | 108 | 86 | 140 |
| | | N50 (bp) | 206,044 | 190,808 | 212,979 | 190,412 | 222,051 | 184,466 |
| | | #mismatches / 100 Kb | 3.07 | 1.78 | 67.11 | 1.96 | 287.37 | 2.03 |
| | | #indels / 100 Kb | 0.16 | 0.06 | 2.03 | 0.13 | 262.26 | 0.11 |
| Long read-only | Conservative | Total length (bp) | 5,326,801 | 5,260,741 | 4,806,912 | 6,307,464 | 5,539,158 | 5,236,419 |
| | | Number of contigs | 6 | 3 | 3 | 22 | 3 | 7 |
| | | N50 (bp) | 5,126,679 | 5,039,909 | 4,797,749 | 5,073,008 | 5,286,804 | 4,924,536 |
| | | #mismatches / 100 Kb | 276.23 | 241.39 | 2,772.51 | 344.5 | 0 | 332.79 |
| | | #indels / 100 Kb | 252.29 | 264.7 | 265 | 306.03 | 0 | 289.71 |
| | Normal | Total length (bp) | 5,326,801 | 5,260,741 | 4,806,912 | 6,307,464 | 5,539,158 | 5,236,419 |
| | | Number of contigs | 6 | 3 | 3 | 22 | 3 | 7 |
| | | N50 (bp) | 5,126,679 | 5,039,909 | 4,797,749 | 5,073,008 | 5,286,804 | 4,924,536 |
| | | #mismatches / 100 Kb | 276.23 | 241.39 | 2772.51 | 344.5 | 0 | 332.79 |
| | | #indels / 100 Kb | 252.29 | 264.7 | 265 | 306.03 | 0 | 289.71 |
| | Bold | Total length (bp) | 5,326,801 | 5,260,741 | 4,806,912 | 6,307,464 | 5,539,158 | 5,236,419 |
| | | Number of contigs | 6 | 3 | 3 | 22 | 2 | 7 |
| | | N50 (bp) | 5,126,679 | 5,039,909 | 4,797,749 | 5,073,008 | 5,286,804 | 4,924,536 |
| | | #mismatches / 100 Kb | 276.23 | 241.39 | 2772.51 | 344.5 | 0 | 332.79 |
| | | #indels / 100 Kb | 252.29 | 264.7 | 265 | 306.03 | 0 | 289.71 |
| Hybrid | Conservative | Total length (bp) | 5,272,824 | 5,275,251 | 5,215,332 | 5,323,049 | 5,055,625 | 5,492,517 |
| | | Number of contigs | 52 | 6 | 191 | 34 | 51 | 107 |
| | | N50 (bp) | 1,444,640 | 5,048,264 | 426,378 | 2,673,977 | 1,423,856 | 749,550 |
| | | #mismatches / 100 Kb | 1.63 | 242.24 | 2,764.2 | 2.04 | 285.57 | 3.7 |
| | | #indels / 100 Kb | 0.32 | 265.38 | 263.44 | 0.09 | 263.18 | 0.02 |
| | Normal | Total length (bp) | 5,276,305 | 5,275,251 | 5,291,108 | 5,327,833 | 5,098,966 | 5,516,886 |
| | | Number of contigs | 42 | 6 | 110 | 33 | 44 | 74 |
| | | N50 (bp) | 1,746,191 | 5,048,264 | 72,0730 | 2,675,388 | 1,762,353 | 1,243,293 |
| | | #mismatches / 100 Kb | 1.56 | 242.24 | 44.59 | 2.28 | 284.11 | 1.65 |
| | | #indels / 100 Kb | 0.28 | 265.38 | 4.07 | 0.13 | 266.82 | 0.02 |
| | Bold | Total length (bp) | 5,293,427 | 5,275,251 | 5,267,003 | 5,223,433 | 5,115,410 | 5,550,270 |
| | | Number of contigs | 23 | 6 | 32 | 3 | 22 | 47 |
| | | N50 (bp) | 3,801,465 | 5,048,264 | 1,222,073 | 3,699,451 | 4,958,323 | 1,266,683 |
| | | #mismatches / 100 Kb | 271.47 | 242.24 | 2,770.38 | 321.64 | 283.97 | 296.99 |
| | | #indels / 100 Kb | 252.55 | 265.38 | 264.11 | 268.47 | 268.27 | 268.29 |

Supplementary Table 4a.

List of genes (with count # indicated by "_") in the plasmid contigs of VREC1013, VRES1160 and VREC1428. The *bla*$_{CTX-M}$ (*bla*), *traI* and *traD* genes are in bold. Only isolate VREC1428 had *traI* and *traD* genes indicating conjugative capacity. VREC0693, VRES0739 and VREC1073 contigs did have *tra* genes.

| VREC1428 | VREC1013 | VRES1160 |
|---|---|---|
| *pemI* | *xerD_1* | ***bla_1*** |
| *pemK* | *ccdA* | *tmrB* |
| ***bla*** | *vapC_1* | *cat* |
| *nqrC* | *vapC_2* | ***bla_2*** |
| *lolD* | *kdgT_1* | *aacA4* |
| *agp* | *kdgT_2* | *tetA_1* |
| *hemR* | *ridA* | *tetA_2* |
| *repB_1* | *yagE* | *pinR* |
| *repB_2* | *ugpA* | *sopB_1* |
| *mmuM* | *cpdA* | *sopB_2* |
| *rocC* | *tnpA* | *repE* |
| *ccdA* | *yknY* | *ccdB* |
| *ccdB* | *tpd* | *ccdA* |
| *sopB_1* | *xerD_2* | *pifC_1* |
| *sopB_2* | *dhfrI_1* | *pifC_2* |
| *klcA* | *ant1_1* | *pifC_3* |
| ***traD_1*** | *folP* | *repB_1* |
| ***traD_2*** | *srpC* | *repB_2* |
| ***traD_3*** | ***bla*** | *xerD* |
| ***traD_4*** | *xerD_3* | *chrA* |
| ***traD_5*** | *xerC* | *folP* |
| ***traD_6*** | *dhfrI_2* | *mdtJ* |
| ***traI_1*** | *ant1_2* | *xerC* |
| ***traI_2*** | *umuC* | *tetA_3* |
| ***traI_3*** | *lexA* | *tetR* |
| ***traI_4*** | *klcA* | *neo* |
| ***traI_5*** | | *tnpR* |
| ***traI_6*** | | |
| ***traI_7*** | | |
| *finO* | | |

Supplementary Table 4b.

Protein products encoded by the genes found in the plasmid of strains VREC1013, VRES1160 and VREC1428 (Figure 2).

| Gene | Protein product |
|---|---|
| *agp* | Glucose-1-phosphatase |
| *ccDA* | Antitoxin (Plasmid maintenance) |
| *chrA* | Response regulator |
| *finO* | Fertility inhibition protein |
| *folP* | Dihydropteroate synthase |
| *hemR* | Hemin TonB-dependent receptor |
| *klcA* | Antirestriction protein |
| *lolD* | Lipoprotein-releasing system ATP-binding |
| *neo* | Aminoglycoside 3'-phosphotransferase |
| *nqrC* | Na(+)-translocating NADH-quinone reductase |
| *pemK* | mRNA interferase |
| *pifC* | Transcriptional repressor protein |
| *pinR* | Serine recombinase protein |
| *repB* | Replication protein |
| *repE* | Replication initiation protein |
| *rocC* | Amino-acid permease |
| *sopB* | Inositol phosphate phosphatase |
| *tetA* | Tetracycline resistance protein |
| *tmrB* | Tunicamycin resistance protein |
| *tnpR* | Transposon gamma-delta resolvase |
| *traD* | Coupling protein |
| *traI* | Multifunctional conjugation protein |
| *xerC* | Tyrosine recombinase protein |
| *xerD* | Tyrosine recombinase protein |

Supplementary Table 5. Long contigs (lengths > 20 Kb) classified as plasmid-derived after alignment with the database of 10,892 complete plasmids [37] to identify the most similar plasmids using BLAST matches spanning more than one gene (match length > 1,000 bp) with a sequence ID threshold of 95%. This showed the most similar plasmids were isolates were spread across *Enterobacteriaceae* for five and one was in Gammaproteobacteria *Shewanella bicestrii* (VRES1160's plasmid), and that relatively high matching levels were detected for VREC0693's and VREC1428's plasmids, but not for VREC1013, VREC1073 nor VRES1160.

| **Strain** | **Length (bp)** | ***bla*$_{CTX-M}$ allele** | **Plasmid type** | **Accession** | **Species** | **Strain name** | **Plasmid name** | **Plasmid length** | **Sum matches > 1,000 bp** |
|---|---|---|---|---|---|---|---|---|---|
| VRES1160 | 61,934 | 15 | IncFIA | NZ_CP022359 | *Shewanella bicestrii* | JAB-1 | pSHE-CTX-M | 193,338 | 30,225 |
| VREC0693 | 132,042 | None | IncFIB | NZ_CP015131 | *Klebsiella pneumoniae* | Kpn555 | pKPN-7c3 | 142,858 | 98,455 |
| VREC0693 | 88,790 | None | IncB | NC_017675 | *Salmonella enterica* | ST4/74 | plasmid TY474p2 | 86,908 | 77,323 |
| VREC1013 | 89,945 | 15 | IncFII | NZ_CP010225 | *Escherichia coli* | M19 | plasmid D | 11,321 | 5,925 |
| VREC1073 | 156,298 | None | IncFIA | NZ_CP015503 | *Klebsiella pneumoniae* | SKGH01 | plasmid unnamed 3 | 84,941 | 39,187 |
| VREC1428 | 92,750 | 27 | IncFIA | NZ_CP022458 | *Shigella sonnei* | 2015C-3566 | plasmid unnamed1 | 55,820 | 53,995 |
